# 3D bone printing via primed differentiation of stem cells with ultrasound (3DBonUS)

**DOI:** 10.1101/2025.11.04.686554

**Authors:** Martina Marcotulli, Chiara Patuto, Chiara Scognamiglio, Federico Serpe, Lucia Iafrate, Efsun Senturk, Sara Keller, Davide De Grandi, Biagio Palmisano, Alessandro Corsi, Mara Riminucci, Giancarlo Ruocco, Dario Carugo, Gianluca Cidonio

## Abstract

Bone disorders and skeletal defects represent a significant clinical challenge. Tissue engineering and regenerative medicine (TERM) strategies using 3D bioprinting have emerged as promising alternatives, but still limited by the inability of directing stem cell differentiation in a controlled and reproducible manner. Advancing beyond this, we are proposing 3D bone printing via primed differentiation of stem cells with ultrasound (referred to here as ‘3DBonUS’). This approach synergistically integrates low-intensity pulsed ultrasound (LIPUS) with a microfluidic-assisted 3D bioprinting platform enabling the biophysical stimulation of human bone marrow stromal cells (HBMSCs) during extrusion, promoting osteogenic differentiation without the need for post-fabrication treatments. Moreover, the incorporation of microbubbles enhanced the effects of LIPUS by amplifying mechanical signals at the cellular level. 3DBonUS was found to significantly upregulate key osteogenic markers (RUNX-2, ALP, COL1A1, BMP-2, OCN, OPN) as confirmed by immunofluorescence and RT-qPCR analysis. Furthermore, the LIPUS-treated constructs showed a significant increase in alkaline phosphatase activity and calcium deposition, indicating enhanced mineralisation. The 3DBonUS strategy represents a new modality in skeletal biofabrication, harnessing targeted minimally-invasive mechanical stimulation, with potential for manufacturing scalability and clinical application. Future studies will aim to validate 3DBonUS in vivo to assess the ultimate regenerative potential with enhanced osteogenic properties.

## 1. Introduction

Bone injuries and defects represent a significant global health burden, often resulting from trauma, neoplasia and infection^1,2^. These conditions are associated with significant morbidity, including chronic pain, functional impairment and recurrent hospitalisation. Autologous bone grafting is the current gold standard treatment. However, this approach is limited by donor tissue availability, donor site morbidity and prolonged operative times^3^. These limitations are alleviated by a number of alternatives such as allografts and xenografts, but these approaches carry risks related to immunogenicity and pathogen transmission^4^. Artificial implants, while offering immediate structural support, often demonstrate suboptimal osseointegration while being typically prone to failure^5^. To overcome the limitations of conventional treatments, tissue engineering and regenerative medicine (TERM) has emerged as a promising therapeutic strategy for skeletal regeneration^6^. Typical approaches involve developing biocompatible and biodegradable scaffolds that support cellular adhesion, proliferation, and differentiation^7^. These scaffolds aim to replicate the complex biomechanical and physiological characteristics of native bone tissue, thereby providing a framework for new tissue formation while undergoing controlled biodegradation^8^. Nevertheless, the efficacy of these methodologies is constrained by the lack of manufacturing approaches to accurately regulate the spatiotemporal signals that govern cell differentiation and tissue maturation.

A notable advance in this area is represented by the advent of 3D bioprinting technology, which facilitates the precise fabrication of cell-embedded scaffolds with unprecedented architectural control^9^. 3D bioprinting typically uses syringe-based printheads to control the deposition of biomaterials and cells in defined patterns^10^. The printheads can be integrated with traditional 3D printers equipped with axial (x, y, z) systems^11^ or incorporated into more advanced robotic arms^12^, offering increased precision and flexibility for patient-specific applications such as targeted tissue regeneration and reconstructive surgery^13^. A key advantage of TERM and 3D bioprinting is the possibility of incorporating human bone marrow stromal cells (HBMSCs), which are inherently capable of differentiating into multiple specialised cell types, including osteoblasts, adipocytes and chondrocytes^14,15^. Indeed, by providing precise biochemical and mechanical stimuli, HBMSCs can be directed towards osteogenic or other relevant lineages^16–18^, enabling the fabrication of implants that are tailored to the specific biological needs of the patient. However, current approaches rely on post-printing stimulation, necessitating additional steps in the long-term to induce osteogenic differentiation following scaffold fabrication. This limitation greatly hinders the real-time control of cell fate and reduces the efficiency of TERM strategies, considerably hindering their clinical translation.

To address this critical limitation, recent advancements have explored the use of external biophysical stimuli, such as low intensity pulsed ultrasound (LIPUS)^19^, to enhance HBMSCs differentiation during scaffold fabrication. LIPUS has shown promise in facilitating bone regeneration through the non-invasive delivery of precise mechanical signals to target cells^20^. Through manipulation of ultrasound-related parameters such as intensity, acoustic pressure and treatment time^21–23^, LIPUS creates a mechanical pressure field that can activate a series of molecular processes, including the release of cytokines, growth factors and adhesion molecules that collectively promote cell proliferation, migration and differentiation^14^. Several studies have shown that LIPUS promotes osteogenic differentiation in 2D models, by upregulating key bone-related genes such as collagen type I alpha 1 (COL1A1), alkaline phosphatase (ALP), osteocalcin (OCN) and bone morphogenetic protein-2 (BMP-2)^24–27^. Furthermore, LIPUS has been shown to promote type-IX collagen accumulation and enhance the proliferation in collagen scaffolds cultivated in a 3D environment with chondrocytes^28^. The osteogenic potential of LIPUS can be significantly enhanced when combined with gas microbubbles (MBs), often creating a synergistic effect that enhances HBMSCs differentiation^29^. When exposed to LIPUS, MBs undergo volumetric oscillations (a phenomenon known as ‘cavitation’), imparting mechanical forces on adjacent cells and promoting cellular activities^30^. Notably, cavitation^31,32^ can transiently permeabilise cell membranes and other biological barriers, facilitating penetration of biologically active compounds^33^.

Despite the advances in 3D bioprinting and LIPUS technologies, no current platform integrates these methods to fabricate a 3D bioprinted construct whilst actively directing cell fate through localised and continuous mechanical stimulation. To address this gap in technology, here we introduce a new platform that achieves 3D bone printing via primed differentiation of stem cells with ultrasound (referred to as ‘3DBonUS’, **Figure 1**). This platform is capable of delivering LIPUS during the 3D bioprinting process, to specifically enhance the osteogenic differentiation of HBMSCs. The incorporation of MBs is intended to further enhance cellular responsiveness to LIPUS and promote more efficient differentiation into bone tissue. The proposed 3DBonUS platform may constitute a paradigm shift in the field of TERM, as it holds potential for *in situ* regulation of cellular behaviour at the time of scaffold fabrication. By combining LIPUS with 3D bioprinting, this research aims to produce functional, implantable scaffolds designed for osteogenic outcomes. The results presented here demonstrate the potential of integrating mechanical stimulation with biofabrication, paving the way for practical and scalable methodologies for the production of personalised and clinically translatable solutions addressing complex skeletal defects.

**Figure 1.**
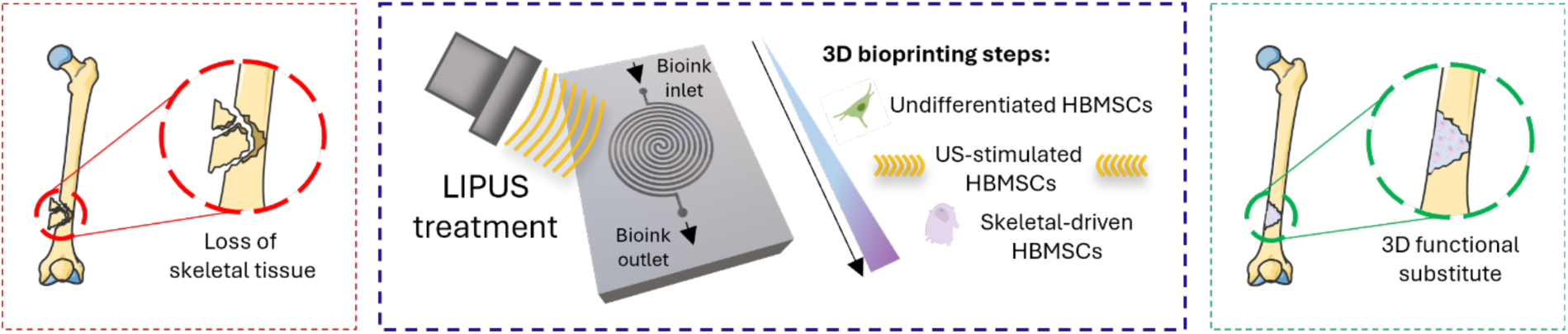
Rationale of 3DBonUS. The process involves the use of a microfluidic printhead designed to drive stem cells to osteogenic differentiation via LIPUS stimulation, enabling the creation of customisable, biodegradable 3D skeletal constructs for bone repair.

## 2. Materials and methods

### Materials

Gelatin from porcine skin (∼300 g Bloom, Type A), Dulbecco′s Phosphate Buffered Saline (PBS), 4-(2-Hydroxyethyl)piperazine-1-ethanesulfonic acid (HEPES), Methacrylic anhydride (MA), 2-(N-Morpholino) ethanesulfonic acid (MES) monohydrate, 2-aminoethyl methacrylate hydrochloride (AEMA), N-hydroxysuccinimide (NHS), 1-[3-(Dimethylamino)-propyl]-3-ethylcarbodiimide methiodide (EDC), 1,2-Distearoyl-sn-glycero-3-phosphocholine (DSPC), Polyethylene glycol 40 stearate (PEG40S), Sodium chloride (NaCl), Glycerol, Lithium phenyl-2,4,6-trimethylbenzoylphosphinate (LAP), Collagenase D, Minimum Essential Medium Eagle – Alpha modification (αMEM), L-Glutamine, Penicillin-Streptomycin, Fetal bovine serum – non-USA origin (FBS), Trypsin-EDTA solution (1x), β-Glycerophosphate disodium salt hydrate (β-GP), 2-Phospho-L-ascorbic acid trisodium salt (AA2P), Dexamethasone, Formaldehyde solution (37% in water), Triton™ X-100, Bovine Serum Albumin (BSA), 4′,6-Diamidino-2-phenylindole dihydrochloride (DAPI), Naphtol AS-MX Phosphate Alkaline, Fast Violet Salt, 3,4-Dihydroxy-9,10-dioxo-2-anthracenesulfonic acid sodium salt (Alizarin Red S), Ammonium hydroxide solution (25%), Latex beads (amine-modified polystyrene, fluorescent yellow-green, 1 µm) and polystyrene-based microparticles (red fluorescent, 10 µm) were purchased from Sigma Aldrich. Calcein (AM, cell-permeant dye), Propidium Iodide, Goat anti-Rabbit IgG (H+L) Cross-Adsorbed Secondary Antibody – Alexa Fluor™ 488, Goat anti-Mouse IgG (H+L) Highly Cross-Adsorbed Secondary Antibody – Alexa Fluor™ 594, Osteopontin Monoclonal Antibody, Osteocalcein Polyclonal Antibody and TRIzol Reagent were purchased from ThermoFisher Scientific. Ethanol absolute anhydrous (EtOH), chloroform and acetone were purchased from Carlo Erba. Alginate (Protanal® GP 1740), SYLGARD™ 184 Silicone Elastomer Kit (PDMS) and Invicta 907 were obtained from FMC BioPolymer, Dow, and Rimas Engineering systems, respectively. Agarose was purchased from Carbosynth. Perfluorobutane (PFB) gas was obtained from F2 Chemicals.

### Synthesis of Gelatin Methacryloyl

Gelatin Methacryloyl (GelMA) was synthesised according to a previously reported method^34,35^. Gelatin (10%) was dissolved in PBS and maintained at 50 °C. After 1 hour, MA (0.8 g per gram of gelatin) was added and the reaction was halted after 3 hours. The solution was then dialysed (1 kDa cut-off) with deionised water (DW) for 5 days. The final GelMA product was frozen at −80 °C, lyophilized (LyoQuest, Teslar), and stored at 4 °C until needed.

### Synthesis of Alginate Methacrylate

Alginate Methacrylate (AlgMA) was synthesised using a previously established protocol^36^. Sodium alginate was dissolved (2% w/v) in 100 ml of 0.05 M MES buffer (0.5 M NaCl, pH 6.5). To activate carboxyl groups, NHS (0.53 g) and EDC (1.75 g) were added to the solution and left to stir for 5 min at room temperature (rt). Subsequently, AEMA (0.76 g) was introduced and the reaction was allowed to proceed with continuous stirring at rt for 24 hours in the dark. The resulting solution was precipitated in 500 ml of cold acetone, collected via vacuum filtration, washed with acetone, and dried under vacuum. The dried AlgMA was rehydrated (1% in DW) and dialysed against DW (3.5 kDa cut-off) for 3 days. Finally, the solution was filtered, freeze-dried, and stored at −20 °C for further use.

### Supporting bath for 3D bioprinting

The supporting bath was prepared according to a procedure reported previously^34,37^. Agarose was dissolved in DW (100 ml) to achieve a concentration of 0.5% w/v. The solution was then autoclaved at 121 °C for 20 min and continuously stirred at 700 RPM for 12 hours. The resulting fluid-gel was stored at rt until further use.

### Manufacturing and characterisation of microbubbles

MBs were synthesised using a previously established protocol with minor modifications^38^. DSPC and PEG40S (9:1 molar ratio) were dissolved in chloroform and allowed to evaporate overnight to form a dry lipid film. The film was rehydrated with PBS to a final lipid/emulsifier concentration of 1.1 mg/ml and stirred at 75 °C for 1 hour. After cooling on ice, PFB gas was introduced and MBs were generated via mechanical agitation using a tissue homogenizer (Precellys® Evolution, Bertin Technologies) operated at 10,000 RPM for 45 seconds. The resulting MB suspension was centrifuged (300 RCF, 5 min, 4 °C) to remove unincorporated lipids. The size distribution and concentration of MBs were analysed using a Multisizer 4e (Beckman Coulter). For optical analysis, MBs were diluted (1:10 in DW), loaded into a hemocytometer (Bright-Line™, Sigma-Aldrich) and imaged at 40x magnification using an optical microscope (DM500, Leica Microsystems GmbH). Image analysis was performed using the Leica’s LAS X Core software. In experiments involving cells, the suspension of MBs was diluted in culture medium to a 50:1 MB-to-cell ratio.

### Fabrication of the piezoelectric transducer

The ultrasonic field was generated using a 647 kHz piezoelectric (PZT) ceramic disc (PZ26, Ferroperm; 18 mm diameter, 3 mm thickness). The upper surface of the PZT disc was separated into two nonconductive areas, by creating a straight engrave with a diamond cutter. The electrical isolation between these two sub-areas was verified with a handheld digital mustimeter (IDM72, RS PRO). Subsequently, silver conductive paint (Silver Conductive Enamel, RS PRO) was applied onto one edge of the PZT disc to create an electrical connection between one sub-area of the upper surface and the bottom surface of the disc. This configuration facilitated coupling of the PZT disc’s bottom surface with the microfluidic device, as electrical wiring could be applied on the upper surface only. Specifically, two crimped cables (JST ZH, 0.05 mm^2^, RS) were separately soldered to the nonconductive sub-areas of the PZT upper surface and were then connected to a BNC cable (Mueller Electric, Female, RS PRO). The bare cables were braided, insulated with heat-shrink tubing (RS PRO), and the transducer was then integrated into the ultrasonic chain for operation.

### Fabrication of microfluidic chips

Two PDMS microfluidic heads were manufactured: i) a spiral chip (channel width: 500 µm, channel height: 250 µm, spiral diameter: 18 mm) for cell stimulation and ii) a printing chip (400 µm inlet channel, Tesla mixer^39,40^ with 6 units). The moulds were designed using CAD software (Autodesk Fusion 360) and later processed using a 3D editor (NAUTA^®^) and a 3D slicing software (FICTOR^®^). For the spiral chip, the geometric design of the microchannel was fabricated with a radius of 250 µm and a total channel length of 244.59 mm. By approximating the spiral channel as a straight semi-cylinder, the following mathematical formulation was employed to calculate the channel volume (**equation 1**):

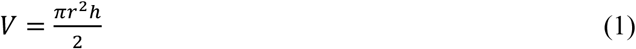

Where *r* is the radius of the channel, *h* is the length of the spiral, and *V* is the volume required to completely fill the channel. A custom-made positive mould was fabricated using an SLA printer (XFAB 3500SD, DWS). After printing, the mould was thoroughly washed with ethanol (EtOH), UV-cured (MeccatroniCore BB Cure) for 10 min (100% power), and then left overnight at 80 °C. Thereafter, PDMS (10:1 w/w base/curing agent) was cast over the mould and cured at 70 °C for 2 hours. The PDMS layer of the spiral chip was plasma-bonded (Harrick Plasma, UK) to a 0.17 mm thick borosilicate glass coverslip (Brand™, Italy) by treatment with oxygen plasma (Plasma Cleaner, Harrick Plasma) for 3 min, while the two halves of the printing chip were bonded together after exposure to plasma for 2 min. For sterile use, EtOH and then PBS were flowed into the microfluidic channels, followed by UV sterilisation (30 min).

### LIPUS exposure

Burst-mode LIPUS signals were generated using a function generator (Tektronix AFG3252C) and a 50-dB power amplifier (1020L RF Power Amplifier, Acquitek). The amplified signal was then transmitted to the 647 kHz PZT transducer. The signal output was constantly monitored with a digital oscilloscope (Tektronix TDS2000C). The acoustic parameters investigated included bursts of 647 kHz sine waves, with varying duty cycles (DC) of 1%, 5%, 15% and 20% (130, 647, 2000 and 2588 cycles, respectively) and a pulse repetition period (PRP) of 20 ms (1 ms pulse duration (PD)). The total LIPUS exposure time during printing was 4 min, which was sufficient for cells to traverse the entire spiral microfluidic channel. Acoustic pressures ranged between 0.2, 0.4, 0.6, 0.8 and 1.0 MPa peak-to-peak (p-p), corresponding to transducer driving voltages of 55, 112, 170, 227 and 288 mV p-p, and input voltages of 25, 50, 75, 100 and 125 V p-p respectively. LIPUS treatment was applied during the bioprinting process, targeting the cells flowing within the spiral microfluidic channel directly above the PZT transducer. For 2D experiments, cells were stimulated similarly to the printing process and were then collected and cultured after LIPUS exposure.

### Acoustic field characterisation

The acoustic pressure field generated by the 647 kHz PZT transducer was characterised using a fibre optic hydrophone (Precision Acoustics, Dorchester) in a degassed deionized water (DW) tank. Experiments were initially conducted to assess the ultrasound field that had travelled through a 0.17 mm thick glass slide, as well as the field that had travelled through both the glass slide and the PDMS layer. The PZT was positioned at the air-water interface of the tank, with the hydrophone placed at 90° with respect to the direction of emission of the ultrasound signal. x-, y-, and z-axis scans were performed to identify the point of maximum pressure (the focal point). Measurements were taken in the near-field (distance of 0.2 mm from the PZT surface), over a 20 mm × 20 mm x-y area. To measure the acoustic field inside the microfluidic device, a PDMS chip with a slit for hydrophone insertion was used. The ultrasound field characterisation was performed at 647 kHz with a 100 V p-p driving voltage (10 cycles), and the transmitting voltage response (TVR) was extracted from the recorded signals using a custom developed script in MATLAB R2023b (The MathWorks, Natick, MA). From this calibration, it was possible to calculate the voltage inputs required to achieve specific acoustic pressure values (**equation 2**):

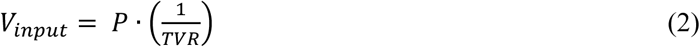

Where *V_input_* is the input voltage (V), *P* is the selected acoustic pressure (Pa), and *TVR* is the hydrophone sensitivity (Pa/V).

### Microbubbles cavitation activity

MBs cavitation activity was assessed via passive cavitation detection (PCD) using a 3.5 MHz focused transducer (V380-SU, Olympus) as a receiver, positioned 10 cm away from the 647 kHz PZT transmitter at a 45° angle. The experimental set-up was submerged in a degassed DW tank, with the PZT transmitter located at the air-water interface. The received signals were amplified using a 5×5×5 gain preamplifier (Panametrics) and digitised with an oscilloscope (DPO7054C, Tektronix) at a sampling frequency of 20 MHz. To reduce background noise, the signals were filtered using a 1.8 MHz high-pass filter (EF509, Thorlabs). MB suspensions at varying dilutions (1:1, 10:1, 50:1 v/v) were conveyed through the spiral microfluidic channel using a syringe pump (Harvard Apparatus) at 6 µl/min. The PCD traces acquired at different acoustic pressures (0.2, 0.4, 0.6, 0.8, 1.0 MPa p-p) were analysed using a custom developed script in MATLAB R2012a (The MathWorks, Natick, MA).

### Biomaterial preparation

The biomaterial was prepared according to a procedure reported previously^41^. Briefly, GelMA (6.5% w/v), AlgMA (5.2% w/v), and LAP (1.3 mM) were mixed in an aqueous solution of NaCl (9 g/L) and HEPES (25 mM) to achieve a final scaffold concentration of 5% w/v GelMA, 4% w/v AlgMA and 1 mM LAP. The solution was stirred for 3 h at 50 °C at 200 rounds per minute (RPM), and LAP was added 10 min before printing to allow chemical cross-linking after printing. Sterilisation was performed by 30 min UV cycles of the powders and filtration (0.22 µm syringe filter, ReliaPrep™, VWR) of the solutions.

### Cell culture and preparation

HBMSCs were isolated and used in accordance with the Declaration of Helsinki and its later amendments with the approval of the Local Ethical Committee (Prot. 0706/2023 - Rif. 7311). The cells were cultured in α-MEM supplemented with 10% FBS, 100 U penicillin, 0.1 mg/ml streptomycin, and 2 mM L-glutamine at 37 °C and 5% CO_2_. Prior to the experiment, cells were thawed and seeded in 175 cm² flasks (5×10^5^ cells/flask, Corning®, NY, USA) with medium replenished twice weekly. At 80% confluency, cells were detached (using Trypsin-EDTA, 1x), pelleted and resuspended at 5.1×10^6^ cells/ml to compensate for dilution in the microfluidic mixer. The final scaffold concentration was 3×10^6^ cells/ml.

### Microfluidic-assisted 3D bioprinting

In the 3DBonUS approach, scaffolds were fabricated using a robotic arm (Rotrics DexArm, Reichelt) with a custom LEGO^®^-based printed support. The cell suspension and biomaterial phase were loaded into 3 ml syringes (Fisherbrand™, Fisher Scientific) and dispensed at controlled flow rates (14 μl/min – biomaterial phase; 6 μl/min – cell suspension phase) via two syringe pumps (Harvard Apparatus, USA). A cylindrical scaffold (7 mm diameter × 3.5 mm height) with 100 µm layer gaps was printed in an agarose fluid gel supporting bath, following g-code instructions via Rotrics Studio. Constructs were then cross-linked *in situ* with a 405 nm UV lamp (5 min) and transferred to well plates (VWR® Multiwell Cell Culture Plates, VWR). The scaffolds were maintained in culture using either i) supplemented α-MEM or ii) osteogenic supplemented α-MEM. The osteogenic medium was prepared by adding 100 μM AA2P, 10 mM β-GP and 10 nM dexamethasone to the supplemented α-MEM.

### Cell viability

To conduct a Live/Dead assay, a solution containing 0.4 μg/ml of Calcein and 0.2 μg/ml of Propidium Iodide was prepared in a FBS-free α-MEM. Cells (for 2D-experiments) or scaffolds (for 3D-experiments) were pre-washed with PBS and incubated for 30 min at 37 °C and 5% CO_2_ in the staining solution. After incubation, the samples were rinsed three times with PBS and examined using a confocal microscope (Olympus IX83) with a 20× objective. Cell viability percentage was analysed quantitatively using Fiji software.

### Immunofluorescence

Cultured samples were fixed in 4% paraformaldehyde (PFA) and permeabilised with 0.1% Triton X-100. For the immunofluorescence (IF) staining of osteocalcin (OCN) and osteopontin (OPN), the samples were incubated in the dark overnight at 4 °C in 1% BSA containing the primary antibodies (anti-Rabbit, 4 μg/ml; anti-Mouse, 2 μg/ml). The secondary antibody staining was performed using OCN antibody (2 μg/ml), OPN antibody (2 μg/ml), and DAPI (1:1000). Finally, the samples were analysed using a confocal microscope (Olympus IX83) with a 40× objective.

### Flow visualisation tests

The flow dynamic field within the spiral microfluidic channel was visualised under LIPUS exposure at varying MB concentrations, using confocal laser scanning microscopy (Eclipse Ti2, Nikon Inc.). Fluorescent yellow-green latex beads (1 µm diameter, 10 µl/ml) were used as flow tracers, while red fluorescent polystyrene microparticles (10 µm diameter, 15 µl/ml) were used to simulate HBMSCs. Suspensions containing MBs at varying dilution ratios (1:1, 10:1, 50:1) and fluorescent beads, were flowed through the microfluidic channel while applying different acoustic pressures (0.2, 0.4, 0.6, 0.8, 1.0 MPa p-p). Images and videos were captured at 10× magnification using NIS-Elements AR software.

### Alkaline Phosphatase staining

The cultured samples were washed twice in PBS, fixed in 95% EtOH and rinsed again in PBS. A DW-based solution containing Naphthol AS-MX Phosphate Alkaline Solution and Fast Red Violet LB Salt was applied over the samples, followed by incubation at 37 °C for 60 min in the dark. Thereafter, the samples were washed three times with PBS and observed using an optical microscope (Zeiss Axiovert) with a 20× objective, with images captured using an integrated camera following standard white balance and fixed brightness/contrast mode. Quantification of ALP activity was carried out using an ALP colorimetric assay kit, and the results were expressed in milli–units per millilitre (mU/ml).

### Alizarin Red S staining

The samples were washed with PBS and fixed in 4% PFA for 1 hour. Thereafter, the cells were stained for 1 hour using a previously prepared Alizarin Red S solution. Briefly, Alizarin Red S (3.5 g) was dissolved in 150 ml of DW, and the pH was adjusted to 4.1-4.3 using 0.5% ammonium hydroxide. The final volume was adjusted to 250 ml while continuously monitoring the pH. Following staining, the samples were washed with DW, and images were acquired using an optical microscope equipped with an integrated camera following standard white balance and fixed brightness/contrast mode. Quantification of Alizarin Red staining intensity was performed using Fiji (ImageJ) software, by analysing n=5 representative regions of interest (ROI) per sample under identical threshold and contrast settings. The mean pixel intensity was normalised to the value of the unstimulated control, and results were expressed as fold change in mineralisation.

### Quantitative reverse transcription polymerase chain reaction

The expression levels of osteogenic differentiation-related genes (**Table 1**) in HBMSCs were assessed using quantitative reverse transcription polymerase chain reaction (RT-qPCR). For 2D experiments, the cells were detached using TRK Lysis Buffer (QIAGEN) and a cell pellet was obtained for RNA extraction. For 3D experiments, the hydrogels were degraded with collagenase D (1 mg/ml) to produce a cell pellet and RNA extraction was carried out using TRIzol reagent. RNA was purified with the RNeasy Mini Kit (QIAGEN) and quantified with a NanoDrop spectrophotometer (NanoDrop 2000c, ThermoFisher Scientific). RNA was reverse-transcribed into complementary DNA (cDNA) using the iScript™ cDNA Synthesis Kit (Bio-Rad). RT-qPCR was performed with iTaq™ Universal SYBR® Green Supermix (Bio-Rad) on a Real-Time PCR System (Applied Biosystems ViiA 7, ThermoFisher Scientific) to quantify gene expression. For both 2D and 3D experiments, a cell pellet from day 0 was used as the control for baseline comparison. Gene expression was analysed using the ΔΔCt method, normalised to the housekeeping gene GAPDH, and presented as fold change (2^-ΔΔCt^) relative to the control sample.

**Table 1.**
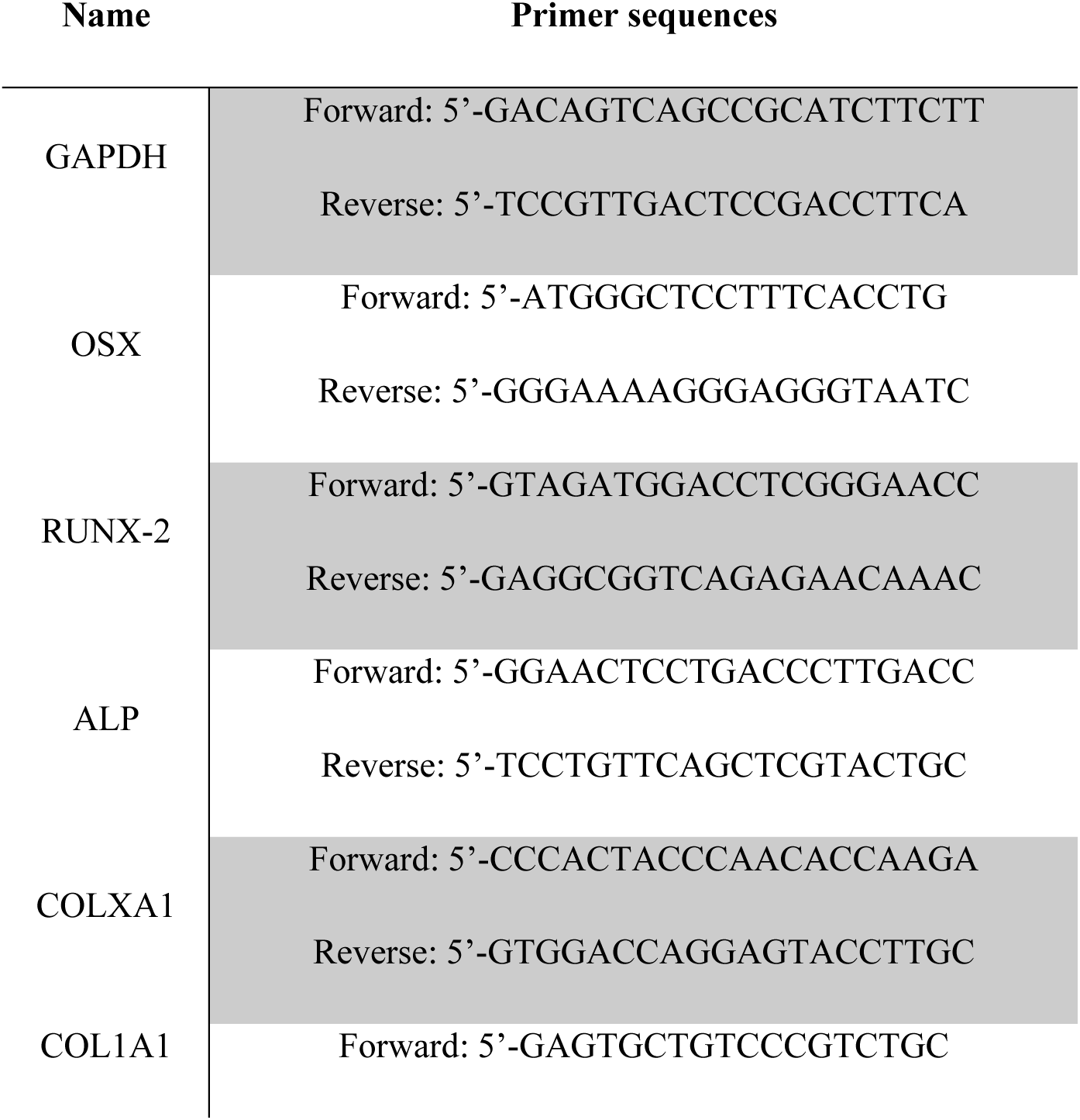
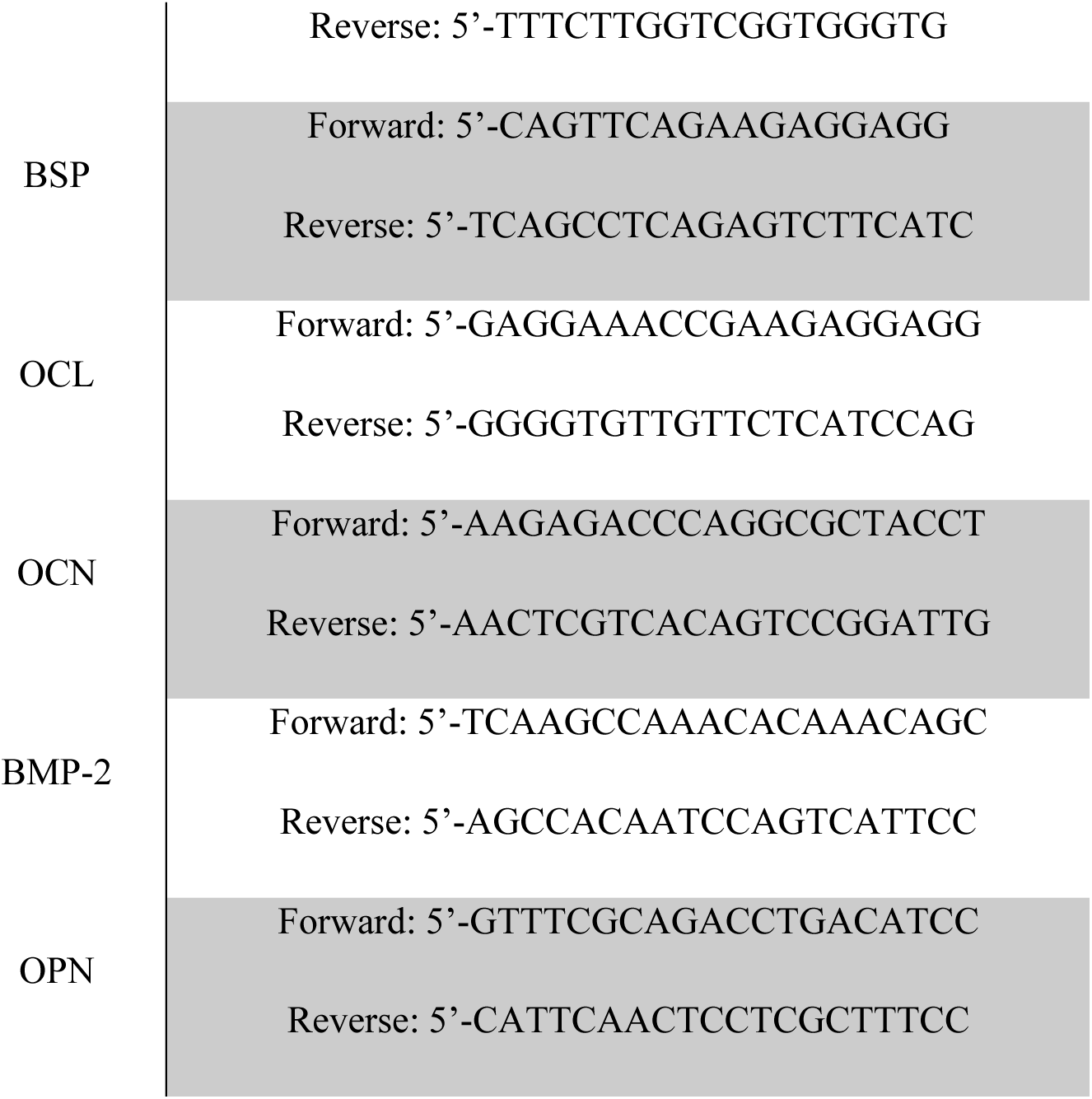
Primer sequences used for RT-qPCR.

### Computational simulation of the acoustic pressure field

To more comprehensively evaluate the acoustic pressure field within the microfluidic device, numerical simulations were conducted using COMSOL Multiphysics 5.5. A 3D model of the set-up was created in CAD software (Autodesk Fusion 360). The design of individual components of the set-up (PZT transducer, glass slide, PDMS layer) were saved as STL files and imported into COMSOL for simulation. The “Pressure Acoustics Frequency Domain” module was employed to calculate the total acoustic pressure field, with a 647 kHz LIPUS source applied at the PZT-glass slide interface. The acoustic impedance values assigned to the set-up components were 13.0 MRayls for the glass slide, 1.48 MRayls for water (simulating the cell suspension medium), and 1.05 MRayls for PDMS (representing the microfluidic chip). Since the model does not allow precise quantification of the total acoustic pressure values, the obtained results were normalised to the maximum acoustic pressure value.

### Statistical analysis

Data analysis was carried out using GraphPad Prism software (La Jolla). The D’Agostino-Pearson normality test was employed to check for normal distribution in the datasets. Differences between groups were analysed using one-way and two-way ANOVA, depending on the experimental setup. The sample size (n) is indicated in the corresponding figure caption. Statistical significance was considered for p<0.05.

## 3. Results and Discussion

To the best of the authors’ knowledge, this is the first study reporting on the direct integration of LIPUS and 3D bioprinting, with the aim of achieving real-time priming of HBMSCs toward osteogenic differentiation and thus eliminating the need for prolonged post-printing treatments. Notably, the proposed method could significantly accelerate and streamline the production of functional skeletal constructs. The following sections describe the functional characterisation, in terms of both physical and biological performance, of the 3DBonUS platform.

### 3.1 Characterisation of the acoustic field of LIPUS in the microfluidic channel system

Experimental and numerical analyses provided a comprehensive characterisation of the acoustic pressure distribution within the developed system (**Figure 2a-c**). In the current study, a microfluidic chip was designed with a spiral microchannel and coupled with a PZT element, to allow acoustic stimulation of the cell suspension flowing through the channel. Being the cells located in close proximity to the ultrasound source (i.e., in the near field, with a PZT-to-cell distance of ∼0.17 mm, equivalent to the thickness of the glass), it was important to assess whether the acoustic field within the microfluidic channel was sufficiently homogenous to ensure even stimulation of the cells. The calibration of the PZT (at 647 kHz operating frequency) allowed the characterisation of the LIPUS field both in the presence and absence of the microfluidic device. To compare the experimental data with a numerical model, the total acoustic pressure field within the experimental system was also determined computationally using COMSOL Multiphysics (**Figure 2b** - **Figure S1**). The simulations revealed the presence of pressure maxima (referred to as ‘pressure antinodes’, in red) alternating with pressure minima (referred to as ‘pressure nodes’, in blue) along the z-axis, corresponding to the direction of LIPUS propagation (**Figure S1a, ii - b,iii**), with regions of higher acoustic pressure located in proximity to the glass slide. In the experimental set-up, the HBMSCs were located inside the microfluidic channel – i.e., between the glass slide and the PDMS layer. Results of the numerical simulations revealed the presence of areas of greater acoustic pressure in the x-y plane parallel to the glass slide (**Figure S1b, i-ii**). The spatial distribution of these high-pressure regions along the spiral channel suggests that cells flowing through the microfluidic device undergo relatively uniform acoustic exposure during stimulation.

**Figure 2.**
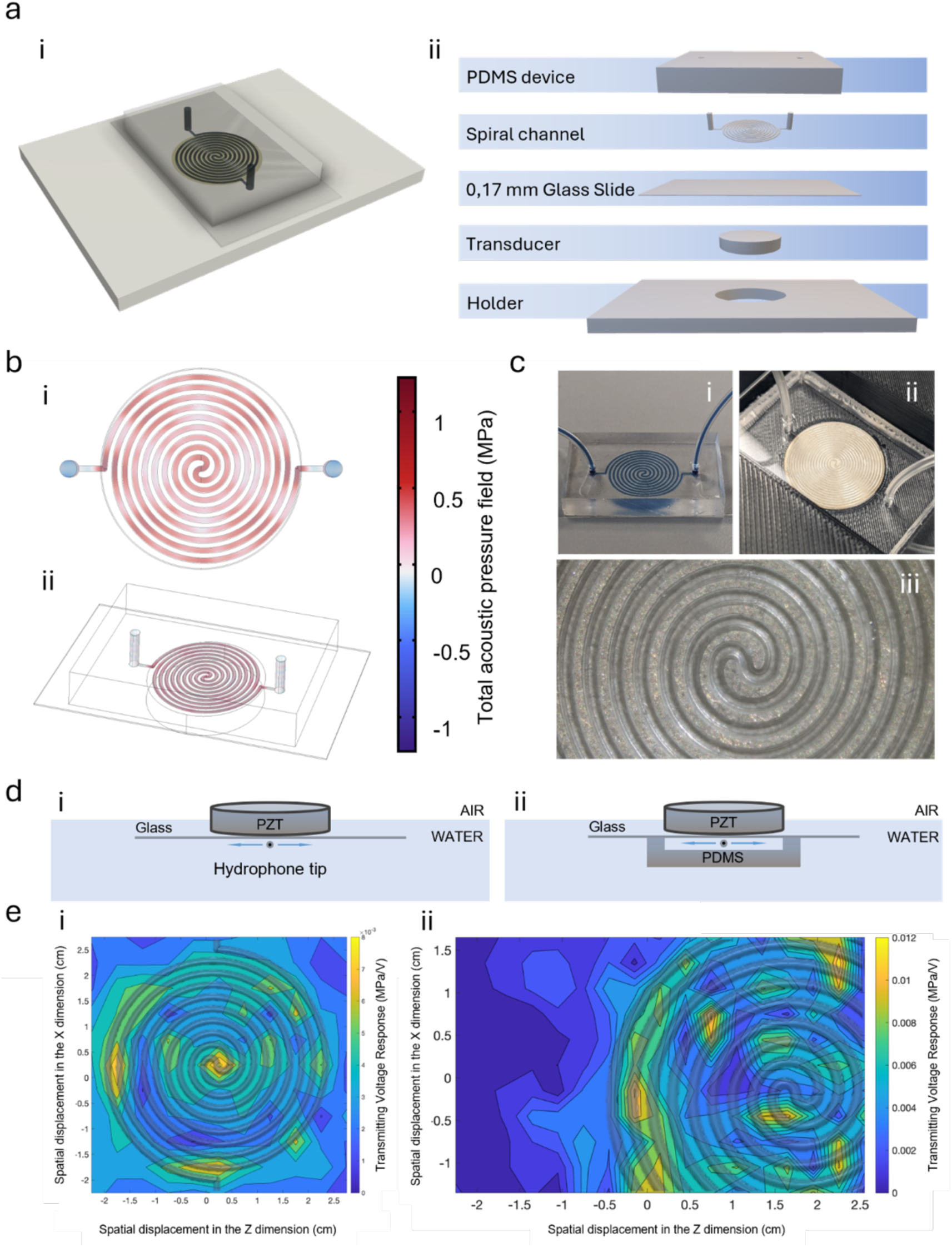
Schematic illustration of the microfluidic device for LIPUS stimulation of cells, and acoustic characterisation of the pressure field emitted by the 647 kHz PZT transducer. a) LIPUS stimulation set-up. a-i) Rendering of the chip observed from the top, and a-ii) rendering of elements forming the LIPUS exposure set-up (from bottom to top: transducer holder, PZT transducer, 0.17 mm glass slide, spiral channel and PDMS layer). b) Rendering in COMSOL of the acoustic device, detailing the total acoustic pressure field. c) Images of the PDMS spiral device. d-i) Configuration of the acoustic calibration set-up of the PZT in the absence of PDMS. The arrows indicate movement in the x-y axis around the hydrophone tip positioned at 90° to the PZT. d-ii) Configuration of the acoustic calibration set-up in the presence of PDMS. The arrows indicate movement in the x-y axis around the hydrophone tip positioned at 90° to the PZT and within an opening created between glass and PDMS. e-i) Merged version of the acoustic pressure field calibration in the presence of the glass slide and the spiral microfluidic channel. e-ii) Merged version of the acoustic pressure field calibration in the presence of PDMS and the spiral microfluidic channel. Acoustic pressure maxima are in yellow, and acoustic pressure minima are in blue.

As illustrated in **Figure S1**, the numerical model provided both the vertical (z-axis) and lateral (x-y plane) distribution of the acoustic pressure field within the device. Considering the device configuration employed, the US wave generated by the PZT element would need to first travel vertically through the 0.17 mm thick glass layer, then through the microfluidic channel containing the cell suspension, and would finally encounter the 5 mm thick air-backed layer of PDMS. Given this layering, it could be anticipated that ultrasound wave reflections may take place at the water-glass interface and, to a lower extent, at the water-PDMS interface, considering differences in acoustic impedance between water (1.48 MRayl), glass (∼13 MRayl)^42^ and PDMS (∼1 MRayl)^43^. Moreover, a thickness of 5 mm for the PDMS layer was specifically chosen to minimise acoustic reflections at the PDMS-air interface. PDMS is known to have a low transmissivity for US waves; therefore, it is anticipated that a relatively thick layer of PDMS would dampen the US wave travelling through it, hence reducing the impact of potential reflections that may take place at the PDMS-air interface^44^. TVR measurements in the near field (with a 0.2 mm distance between the PZT and hydrophone) provided values of 4-8 kPa/V for the PZT with only the glass slide, and of 6-10 kPa/V for the full experimental setup (PZT, glass slide and PDMS) (**Figure 2d**). As shown in **Figure 2e-i**, the acoustic pressure was measured over an area of 20 mm × 20 mm, which corresponded to the location of the spiral microfluidic channel, in order to determine the stimulation conditions of HBMSCs. The experimental calibration, both in the presence and absence of PDMS, revealed the presence of discrete regions of alternating high and low acoustic pressure across the scanned area. The greatest pressure values were detected in the central region of the field (approximately within the inner 4-5 mm), followed by alternating decreases and increases toward the outer region, consistent with the formation of pressure nodes and antinodes previously described. The overall spatial pattern and the peak pressure magnitude obtained from our measurements were found to be similar to those in other compact microfluidic devices^32,45^.

In the presence of PDMS (**Figure 2e-ii**), the acoustic pressure measurements were partially incomplete due to the opening created in the PDMS where the hydrophone was inserted. As was also observed in the absence of PDMS, lateral spatial variations in the acoustic pressure magnitude were detected, likely originating from geometric discontinuities and lateral boundary effects rather than from strong impedance mismatches. Although PDMS has a slightly lower acoustic impedance than water (approximately 1 MRayl vs. 1.40 MRayl), it is generally regarded as acoustically transparent. In the configuration under consideration, the 5 mm PDMS layer functioned principally as a damping medium, thereby reducing the likelihood and impact of potential reflections at the PDMS-air interface. For the LIPUS stimulation experiments, it was important to evaluate the acoustic pressure distribution at the liquid-glass interface. Typically, the PZT-glass coupling effectively behaves as a continuum and the glass mainly acts as a carrier for the US wave. The 0.17 mm glass slide thus allowed minimum distance between the PZT and the cells to be maintained, without significantly affecting the transmission of LIPUS waves into the spiral channel. In order to comprehensively evaluate the acoustic pressure distribution in correspondence to the microfluidic channel, an overlaid representation was created by positioning the pressure field calibration map onto the spiral channel geometry (**Figure 2e**). This composite image was used to identify which regions of the spiral corresponded to areas of higher and lower acoustic pressure.

Notably, the acoustic pressure levels generated during the experiments (0.2 – 1.0 MPa p-p) are within the range commonly used for therapeutic ultrasound applications^46^. The LIPUS parameters chosen for this study were initially derived from values previously employed in the literature. A number of studies have used centre frequencies between 0.5 MHz and 1.5 MHz to stimulate HBMSCs to increase their osteogenic differentiation, with acoustic pressure of approximately 1.0 MPa p-p, and often in burst mode^29,47–49^. Taken all together, both the experimental measurements and the simulations indicate that the assembly of PZT element and microfluidic device is capable of generating a LIPUS field of therapeutically relevant acoustic pressure and frequency over the surface of the spiral channel. Despite the presence of spatial gradients, the experimentally measured acoustic pressure field was consistent with the simulated field, resulting in a relatively homogeneous pressure distribution for the intended application.

### 3.2 Optimised LIPUS stimuli for enhanced osteogenic differentiation of HBMSCs in a 2D environment revealed significant functional effect *in vitro*

Following the characterisation of the ultrasound field in the device, LIPUS stimulation of HBMSCs was first carried out at an acoustic pressure of 1.0 MPa p-p. This pressure was chosen based on its therapeutic relevance^46^ and to avoid overheating observed when using the PZT at pressures higher than 1.2-1.4 MPa p-p. The aim of these initial experiments was to evaluate potential cellular responses to LIPUS in 2D cultures, without the use of biomaterials or 3D bioprinting techniques. Specifically, cells were exposed to LIPUS inside the microfluidic spiral channel and then cultured for up to 21 days. The choice of LIPUS parameters was supported by both the literature and preliminary data from our own experiments. For instance, previous studies^17,50–52^ employing LIPUS for bone and cartilage tissue engineering (TE) applications have commonly used a 20% DC, which was initially adopted in out experimental set-up. However, the DC was later reduced to 5% to mitigate overheating of the PZT transducer. This adjustment was necessary as prolonged use of the PZT at higher DC levels resulted in electrical failure requiring frequent re-soldering. The PRP (20 ms) was also chosen based on consistency with established LIPUS stimulation protocols^53^.

In these experiments (**Figure 3-4**), HBMSCs were stimulated for a total of 4 min, which was determined to be the maximum viable time for effective stimulation prior to extrusion and 3D deposition. The selected flow rate of 6 μl/min within the microfluidic channel was also a critical factor. Lower flow rates led to aggregation of the cells, preventing them from moving uniformly through the channel, whereas above 6 μl/min ensured a steady motion of cells and minimised aggregation. The flow rate value was particularly important as these experiments served as a precursor to our intended use of LIPUS in 3D bioprinting, whereby all cells should ideally be stimulated with a LIPUS field with comparable intensity and duration. The overall goal was to stimulate HBMSCs during the 3D bioprinting process only, creating a cellular scaffold populated with LIPUS-stimulated cells that exhibited accelerated differentiation into bone tissue without the need for post-printing treatments. The design of our system, in which the microfluidic spiral channel was positioned directly above the PZT disc, ensured that all HBMSCs passing through the channel were at a comparable distance from the transducer and received LIPUS stimulation of the same duration. Characterisation of the acoustic field generated by the PZT transducer within the microfluidic channel also confirmed that cells experienced relatively comparable acoustic pressure levels while transiting through the device, thus ensuring similar stimulation conditions across the cell population.

**Figure 3.**
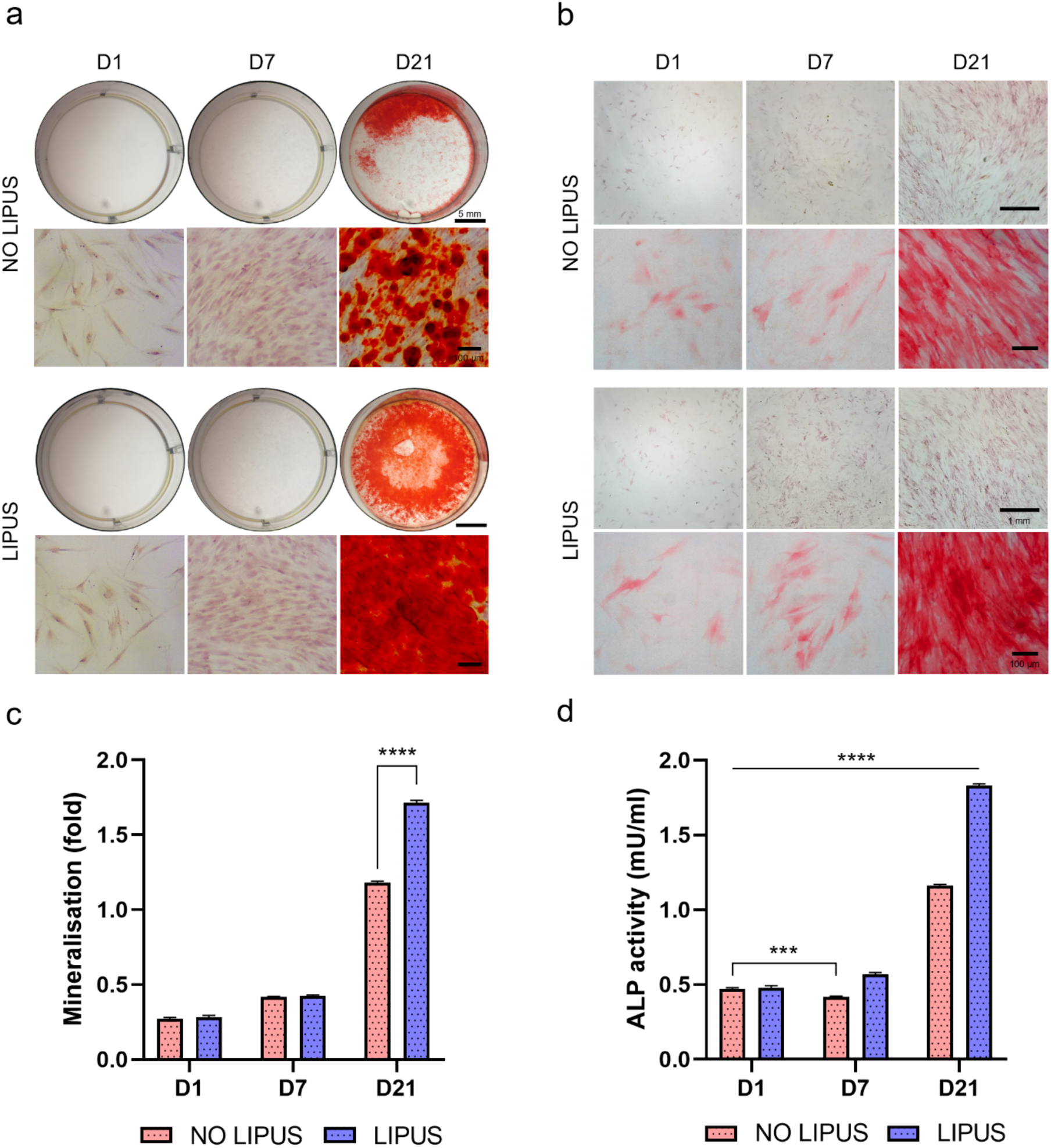
Mineralisation and ALP activity of HBMSCs, with and without LIPUS exposure. a) Ca^2+^ deposits observed at 1-7-21 days after cell culture of non-stimulated and LIPUS-stimulated HBMSCs. Images were taken at both 1x and 20x magnification. b) ALP expression observed at 1-7-21 days after cell culture of non-stimulated and LIPUS-stimulated HBMSCs. Images were taken at both 5x and 20x magnification. c) Mineralisation was quantified in mU/ml. d) ALP activity was quantified in mU/ml. Scale bars: (a) 5 mm, 100 μm, (c) 1 mm, 100 μm. Statistical significance was assessed by two-way ANOVA. Mean ± S.D. n=3, ****p<0.0001, ***p<0.001.

**Figure 4.**
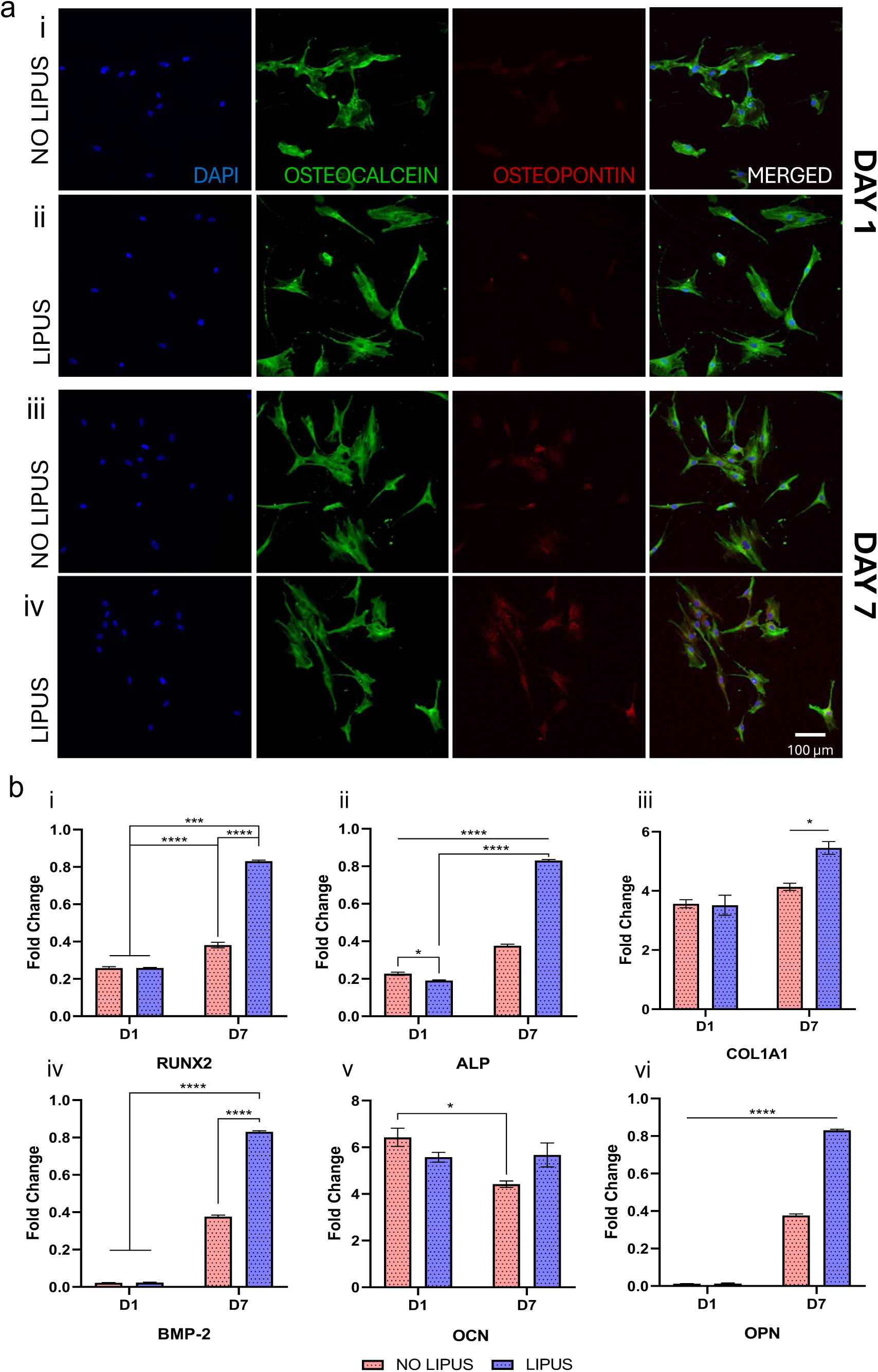
Immunofluorescence images and gene expression of controls and LIPUS-treated HBMSCs. a) Fluorescence images of HBMSCs after 1 (i, ii) and 7 (iii, iv) days of 2D culture showing the nucleus (blue), OCN (green) and OPN (red) at 20x magnification. b) RT-qPCR analysis of HBMSCs after 1 day and 7 days of 2D culture. The expression of genes associated with osteogenic differentiation was examined to observe the effect of LIPUS stimulation. The expression levels of (i) RUNX-2, (ii) ALP, (iii) COL1A1, (iv) BMP-2, (v) OCN and (vi) OPN were significantly higher in samples exposed to LIPUS at day 7 compared to controls. Scale bars: (a) 100 μm. Statistical significance was assessed by two-way ANOVA. Mean ± S.D. n=3, ****p < 0.0001, ***p<0.001, *p<0.05.

Viability assays and staining were performed to assess both cellular vitality and osteogenic differentiation potential of LIPUS-exposed cells compared to unstimulated controls. Live/Dead assay (**Figure S2a,b**) showed that LIPUS exposure did not compromise cell integrity, as the viability of LIPUS-stimulated cells remained high (91.7%±3.3) and comparable to that of unstimulated cells (95.2%±2.3) at day 21, consistent with findings reported in the literature^54,55^. Furthermore, the rate of cell proliferation at day 21 in the LIPUS-stimulated samples was found to be 171.8%±10.8, while the control cells exhibited a proliferation rate of 202.5%±3.6 in comparison to the day 1 data (**Figure S2c**). This reduction in proliferation is consistent with findings in previous studies^56^, which suggest that LIPUS may modulate the cell cycle and promote osteogenic differentiation, at the expense of proliferation. The differentiation of HBMSCs towards skeletal lineage was found to be significant as indicated by a marked increase in calcium deposition (p<0.0001) and ALP activity (p<0.001) by day 21 in LIPUS-stimulated cells, as demonstrated by both Alizarin Red staining for calcium deposits (**Figure 3a**) and ALP staining (**Figure 3b**). Quantification of the intensively stained noduli (**Figure 3c**) and ALP activity (**Figure 3d**) demonstrated a significant difference in the stained area and ultimate functionality of the LIPUS-stimulated groups. These findings were consistent with the literature, reporting increased calcium deposition^57,58^ and ALP expression^24,59^ in HBMSCs exposed to LIPUS stimulation between days 7 and 21.

Immunofluorescence analysis further confirmed that, at day 7, LIPUS-stimulated cells exhibited greater expression of osteogenic markers such as OPN and OCN, in line with the findings by Wang^60^ and Renno^61^ *et al.*, respectively (**Figure 4a**). Following 7 days of culture, the expression of OCN was found to be greater in LIPUS-treated samples than in the controls. While the expression of OPN was less pronounced than OCN, it was nevertheless higher in the LIPUS samples, with the lowest levels observed in the untreated controls. The immunofluorescence investigation was supported by quantitative analysis using RT-qPCR, which showed an upregulation of key osteogenic genes, including RUNX-2, ALP, COL1A1, BMP-2, OCN and OPN, in LIPUS-stimulated cells compared to controls^24,62^ (**Figure 4b**). At day 7, the expression of major osteogenic genes (e.g., RUNX2, ALP, BMP2, OPN) in HBMSCs exposed to LIPUS was significantly (p<0.0001) higher compared to control cells that were not stimulated. Interestingly, COL1A1 and OCN expression was found enhanced (p<0.05) but with a lower degree compared to the unstimulated controls. A study by Costa and co-workers^24^ confirmed that LIPUS stimulation of HBMSCs effectively promotes osteogenic differentiation *in vitro*. This was demonstrated by gene expression analysis of the same genes set observed in our study, suggesting that LIPUS was able to enhance cellular differentiation, partially mimicking the bone microenvironment.

These results suggest that our LIPUS stimulation protocol, characterised by precise control of the acoustic pressure field and timing, successfully promoted HBMSCs osteogenic differentiation in a bidimensional culture micro-environment. The mechanical stimulation provided by the acoustic pressure field likely created an environment suitable for accelerated differentiation into osteogenic lineages, as evidenced by both phenotypic markers and gene expression data. We hypothesise that this mechanism would be enhanced in a 3D environment, by the augmented interaction with extracellular matrix protein and biomaterials. We will also evaluate whether the addition of ultrasound-responsive microbubbles (MBs) would enhance and further modulate HBMSCs response, by amplifying the mechanical effects of ultrasound stimulation.

Overall, in contrast to conventional methods where cells were stimulated for extended periods of time (between 5 to 20 minutes per day, for 7 to 21 days)^63–66^, our 3DBonUS approach focused on a single, controlled LIPUS-exposure event right before the 3D bioprinting process. This strategy not only simplifies the protocol of cell differentiation but also allows for precise temporal control of LIPUS exposure.

### 3.3 Augmenting the functional effect of US on HBMSCs through the inclusion of gas microbubbles

We hypothesised that the addition of MBs to the system would enhance the cellular response of HBMSCs during LIPUS stimulation. MBs are known to amplify cellular activities due to their ability to volumetrically oscillate (a phenomenon known as cavitation) and generate mechanical effects on nearby biological barriers (including cell membranes) when exposed to ultrasound waves^67,68^. To integrate this concept into our established 2D model for LIPUS cellular stimulation, we introduced MBs consisting of a perfluorobutane gas core stabilised by a DSPC/PEG40S shell.

The MBs produced had an initial concentration of 1×10^10^ MBs/ml with an average diameter of 0.68±1.67 µm (**Figure 5a**). After washing and filtration, the concentration decreased to 1×10^9^ MBs/ml, while the average diameter increased to 1.19±1.55 µm (**Figure 5b**). This shift in size and concentration was expected, as the filtration process removed supramolecular lipid aggregates and selectively retained fully formed MBs^69^. To assess how MBs responded to ultrasound stimulation, a PCD test was conducted to evaluate the frequency content of the signal generated by MBs upon LIPUS stimulation. At the selected acoustic pressure of 0.2 MPa p-p, no harmonics beyond the fundamental frequency of 647 kHz were observed (**Figure 5c,i**). However, at 0.6 MPa p-p, three harmonics appeared at 1.29 MHz, 1.94 MHz, and 2.59 MHz (**Figure 5c,ii**). When the pressure was increased further to 1.0 MPa p-p, a fourth harmonic at 3.23 MHz was detected (**Figure 5c,iii**). These findings indicate that increasing the acoustic pressure led to more enhanced MB oscillation, evidenced by the presence of additional harmonics in the signal emitted by the cavitating MBs and supported by previous studies^30^.

**Figure 5.**
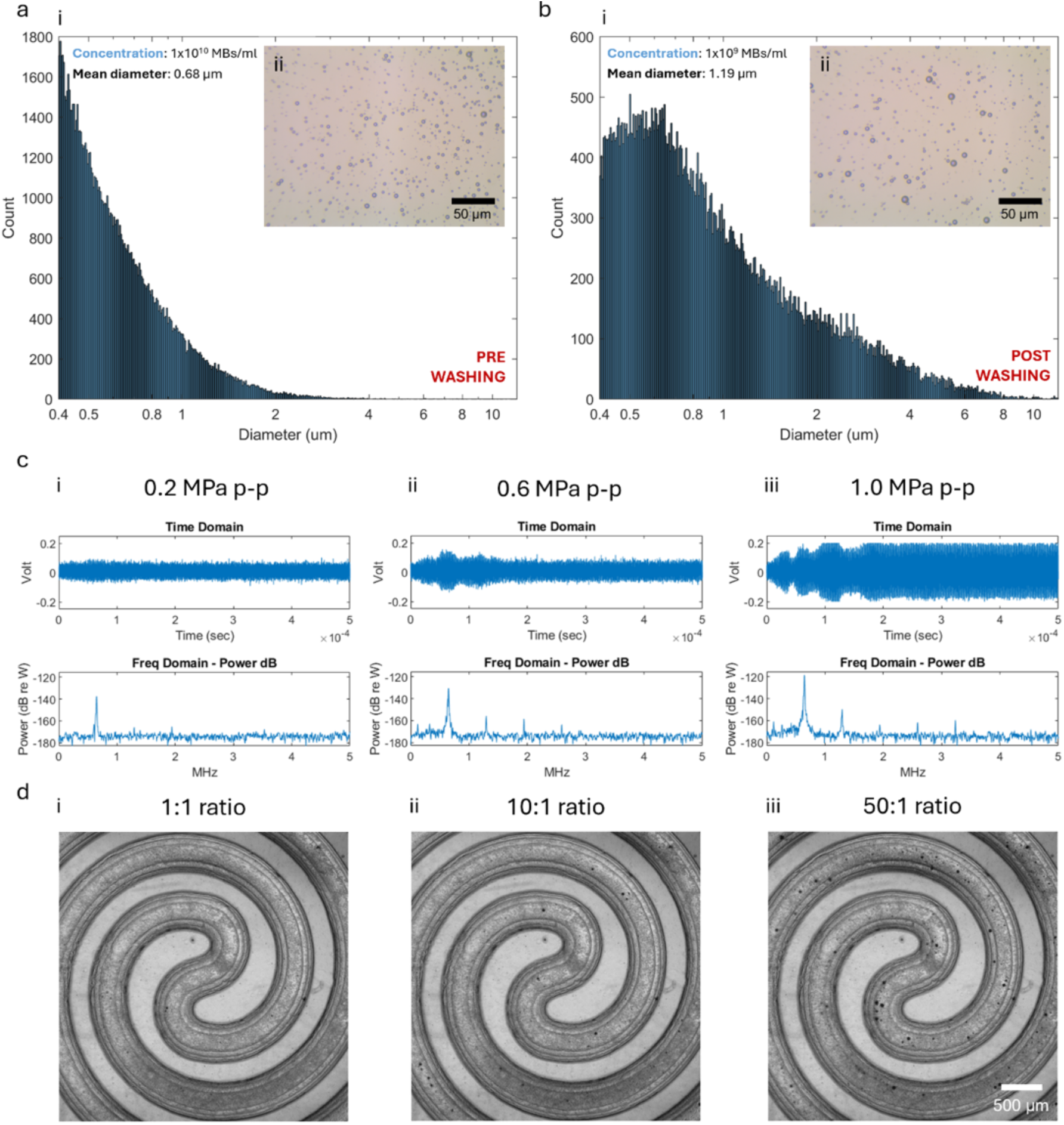
Characterisation of MBs size and cavitation behaviour. a) MBs size distribution in µm a-i) before washing, with a-ii) showing micrographs of the MBs obtained with a 40x objective. b) MBs size distribution in µm b-i) after washing, with b-ii) showing micrographs of the MBs obtained with a 40x objective. c) Time domain and frequency domain obtained from PCD measurements performed at acoustic pressures of c-i) 0.2 MPa p-p, c-ii) 0.6 MPa p-p and c-iii) 1.0 MPa p-p. The time domain plots display the voltage amplitude (V) of the acoustic emission recorded by the receiving transducer as a function of time (sec). The corresponding frequency domain spectra show the acoustic power (dB re W) as a function of frequency (MHz). From left to right, an increase in voltage in the time domain spectrum was observed, while in the frequency domain spectrum harmonics after the fundamental frequency were detected (zero, three and four harmonics in the spectra obtained at 0.2 MPa p-p, 0.6 MPa p-p and 1.0 MPa p-p, respectively). d) Brightfield microscopy images of the microfluidic channel filled with MB-to-cell ratios of d-i) 1:1, d-ii) 10:1 and d-iii) 50:1, respectively. Scale bars are: (a-ii) 50 µm, (b-ii) 50 µm, and (d) 500 µm.

The relatively weak signal intensity and the lack of broadband content suggest that stable cavitation, rather than inertial cavitation^70^, was the dominant cavitation mode in the system. Stable cavitation, characterised by sustained and repeated MB oscillations (and the absence of violent MB collapse), may be preferable for promoting cellular responses whilst minimising cellular damage^71^. Furthermore, at acoustic pressures of 0.6 MPa p-p and 1.0 MPa p-p, sub-harmonics were observed. Sojahrood and colleagues^72^ associated this phenomenon with the large size dispersity of MB populations and potential coalescence events occurring during ultrasound exposure, which can induce non-linear radial oscillations of the MBs. These oscillations are responsible for the generation of sub-harmonic frequencies observed in the detected acoustic spectrum. The 50:1 MB-to-cell ratio was selected in the LIPUS stimulation experiments (**Figure 5d**) based on both prior studies using similar values^73^ and our own experiments, which showed that this MB concentration could fill the spiral channel used. Xu and co-workers^74^ investigated higher concentrations of MBs (100:1 MB-to-cell ratio) and reported that MB-mediated US therapy provided remarkable improvements in cellular targeting and interactions. Lower ratios in the current study, such as 1:1 and 10:1, resulted in insufficient MB numbers in the microfluidic channel, which likely limited MB oscillation under LIPUS exposure. Although this was not directly confirmed by PCD analysis, the reduced acoustic response observed at these ratios supports this interpretation. Following the characterisation of MB response under LIPUS exposure, a novel printing head was developed, capable of simultaneously delivering biomaterials and LIPUS stimulation to HBMSCs in the presence of MBs.

### 3.4 3DBonUS bioprinting with a dual-chip microfluidic system for LIPUS stimulation and simultaneous bone-guided deposition

The characterisation of LIPUS parameters tailored for stem cells stimulation, laid the foundation for subsequent studies evaluating whether the use of the 3DBonUS microfluidic strategy could enhance the osteogenic differentiation of HBMSCs. By exploiting the potential of LIPUS, efforts were made to improve the efficiency and consistency of the differentiation process. To this end, a dual-chip microfluidic system was developed to facilitate both LIPUS exposure and 3D bioprinting, ensuring a streamlined workflow for the preparation and printing of scaffolds conducive to osteogenesis (**Figure 6a**).

**Figure 6.**
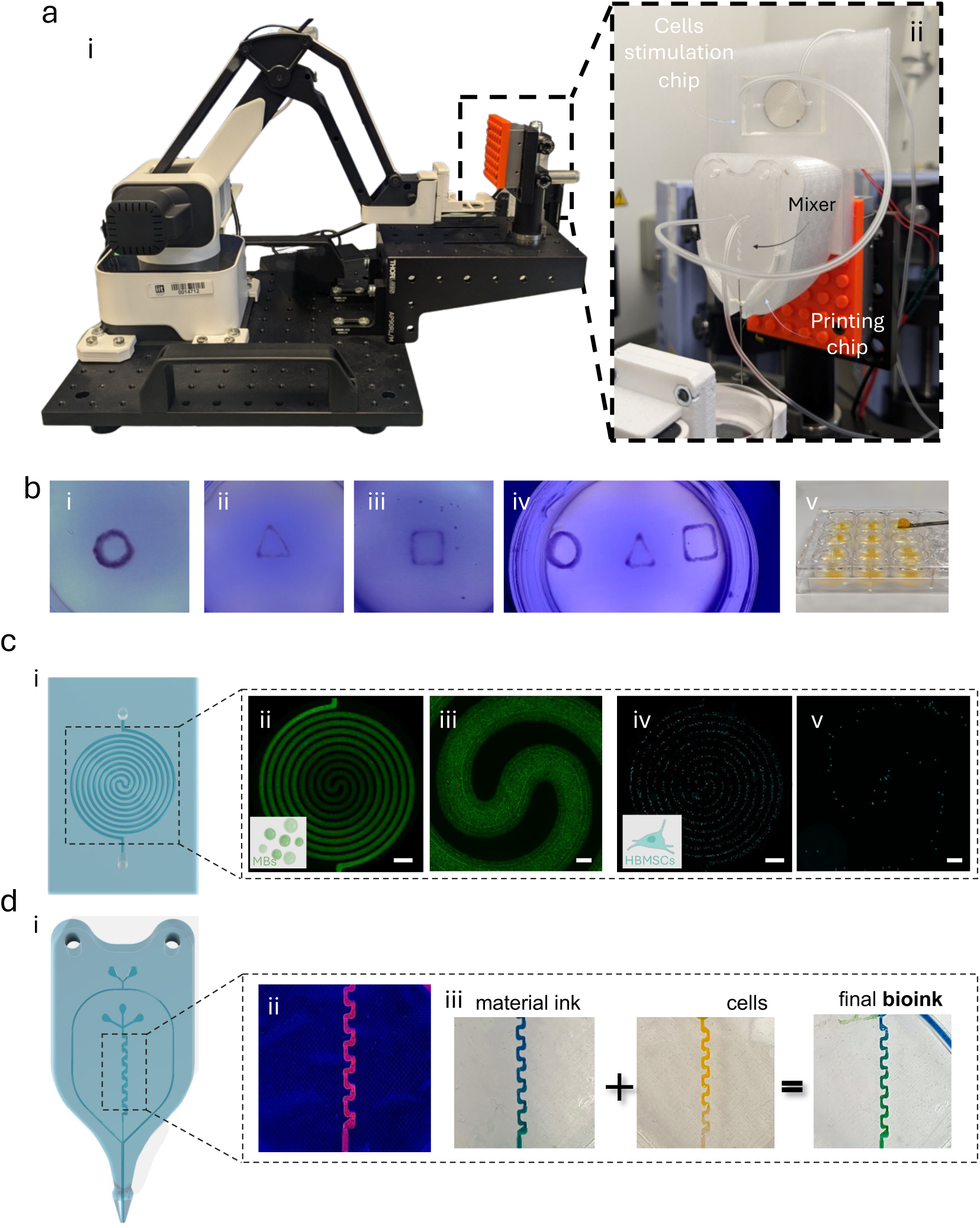
Schematic illustration of the 3DBonUS stimulation set-up. a) Complete set-up for LIPUS stimulation. a-i) Robotic arm for 3D bioprinting with LEGO^®^ support for microfluidic head adhesion. b-ii) 3D support designed using Autodesk Fusion 360 and visualisation of the two microfluidic heads. b) Examples of scaffolds printed using the 3DBonUS method. c) Flow visualisation test performed using c-i, c-ii) 1 µm beads (green) to simulate MBs and c-iii, c-iv) 10 µm beads (cyan) to simulate HBMSCs. d) Flow visualisation in the micro-mixer using d-ii) rhodamine B to simulate the flow, d-iii) blue and yellow dyes to simulate material ink and cell suspension, respectively. Scale bar: (c,i – c,iii) 250 μm, (c,ii – c,iv) 2000 μm.

The first microfluidic chip was designed with a spiral microchannel to allow uniform stimulation of the cells and MBs suspension. In the 3DBonUS system, a flow rate of 6 µl/min was selected, corresponding to a 4-minute residence time (and therefore stimulation time by LIPUS) within the spiral microchannel. This exposure time was selected based on experimental verification and findings reported in the literature, supporting its suitability for inducing osteogenic effects. Following the stimulation of cells, the chip content could be conveyed into the second chip, which is dedicated to the 3D bioprinting process (**Figure 6a**). The printing chip was equipped with two inlets: one for the cell suspension from the first chip and another for the biomaterial solution, which consisted of GelMA and AlgMA in a NaCl + HBSS buffer. A Tesla mixer within the chip ensured homogeneous mixing of the cell-laden bioink prior to the bioprinting step^39^. The first chip (bonded to a glass substrate) required direct contact with the PZT transducer, while the second chip (consisting of two PDMS layers) facilitated precise bioprinting. This modular approach mitigated the complexity of combining LIPUS exposure and biomaterial manipulation within a single chip and addressed the core objective of the study: effectively stimulating cells during printing to reduce experimental complexity.

In terms of biomaterial selection, AlgMA was chosen over traditional alginate due to its superior compatibility and ease of crosslinking *via* photopolymerisation, following deposition in an agarose suspension bath. Notably, conventional alginate necessitates the use of CaCl_2_ for cross-linking following the printing process, greatly limiting post-printing applications when used in combination with support bath printing^75^. Moreover, the methacrylated inks employed in this study (GelMA and AlgMA), allowed UV light to be used for cross-linking within the agarose phase, providing greater control compared to in-air extrusion over the architectural construct. This was further supported by previous experiments^41^, where both materials showed favourable performance in generating water-in-water emulsions, resulting in scaffolds with uniform porosity. By exploiting the physical and chemical cross-linking properties of these materials, a stable 3D construct suitable for osteogenic differentiation was achieved. Flow rate calibration was critical for controlling cell and material concentrations within the final printed scaffold (**Figure 6b**). Specifically, the cell suspension and biomaterial solution were driven at 20 µl/min during printing. This total flow rate used was inspired by previous work^34^ where the same value was employed to print similar structures in agarose suspension. Given the dilution effects inherent in the system, the initial concentrations of cells and biomaterials were adjusted to achieve a final composition of 5% GelMA, 4% AlgMA, 1 mM LAP and a cell concentration of 3 million cells/ml in the printed scaffold. Specifically, considering the ratio between flow rates of the two phases, the initial concentration of the cell suspension was increased by 70%, corresponding to 5.1 million cells/ml, while the biomaterial solution was increased by 30% (6.5% w/v GelMA, 5.2% w/v AlgMA, 1.3 mM LAP) to maintain the final scaffold composition. These adjustments ensured that the scaffold retained the desired mechanical and biological properties for effective osteogenesis after printing. Flow visualisation tests were performed using beads of different size to simulate cells (10 µm) and MBs (1 µm) within the spiral microchannel (**Figure 6c**). In the printing mixer chip (**Figure 6d**), coloured inks were used instead: blue dye represented the biomaterial ink, and yellow dye represented the cell suspension. This allowed the mixing behaviour to be visualised prior to printing. The results confirmed that the microfluidic assembly produced steady laminar flow within the spiral channel. Both 1 µm and 10 µm microspheres followed well-defined streamlines, showing no evidence of aggregation or turbulence. These results verified that the intended flow dynamics, characterised by steady laminar flow, were achieved and that particles of comparable size to cells and MBs were present and uniformly distributed throughout the bioprinting process. This validation step was critical to ensure that the cells were evenly distributed within the scaffold, which is essential for consistent cellular differentiation. The 3DBonUS approach introduces an unprecedented integration of LIPUS stimulation with microfluidic-assisted 3D bioprinting, which differs from previous methods that have primarily used US for tasks such as organising cells into three-dimensional constructs (e.g. US-assisted bioprinting^76^). While US-assisted bioprinting generally relies on external acoustic fields applied post-deposition or in separate chambers to manipulate cell spatial distribution^76^, the developed 3DBonUS method integrates US directly within the microfluidic path during 3D bioprinting to achieve spatially and temporally controlled cellular stimulation. Thus, LIPUS was applied for the first time to a microfluidic system to stimulate HBMSCs in a single-step and continuous-flow modality, rather than relying on multiple daily stimulations over an extended period^63–66^. This innovation simplifies the experimental protocol and provides a more practical and scalable solution for accelerating osteogenic differentiation.

### 3.5 Combined effects of LIPUS and MBs on 3D scaffolds foster osteogenic differentiation

Enhanced osteogenesis was observed, highlighting the potential of integrating LIPUS with microfluidic-assisted 3D bioprinting for bone regeneration purposes. Following the combination of 3D bioprinting and LIPUS, additional analyses were performed to complement those previously carried out in 2D environments. The 3D bioprinted bone-guided scaffolds were cultured for up to 7 days to observe early-stage differentiation, a timeframe chosen due to the inclusion of MBs, which were hypothesised to elicit a more pronounced cellular response compared to 2D experiments. Unstimulated control samples were compared with those stimulated with LIPUS and LIPUS + MBs, to determine whether MBs enhanced the cellular response of HBMSCs.

Cell viability within the GelMA and AlgMA hydrogels was first assessed (**Figure S3a,b**). Previous experiments^41^ showed that HBMSCs maintained high viability in 5% GelMA and 4% Alginate, reaching 91.4%±6.8 on day 7. Consistent with previous results, the present study showed sustained high viability in all three experimental conditions. On day 7, control samples showed 94.1±1.5 % viability, LIPUS-stimulated samples 94.3±1.0 %, and LIPUS+MBs samples 93.3±2.7 %. These results confirmed that LIPUS stimulation did not adversely affect cell viability and corroborated the findings of Jin and co-workers^77^, who reported that mechanical stimulation via LIPUS maintained high cell viability and even enhanced cell proliferation in stimulated scaffolds. Similarly, the rate of cell proliferation at day 7 indicated a comparable increase under all conditions. The proliferation percentage was 58.7%±8.4 for the control samples, 57.1%±8.1 for the stimulated cells and 33.3%±5.4 for the stimulated samples in the presence of MBs, when compared to the initial day 1 data (**Figure S3c**). Calcium deposition by Alizarin Red staining was not performed in the 3D environment due to the high absorption capacity of the porous biomaterial, which prevented accurate detection of calcium deposits and may have led to erroneous conclusions. This problem has been noted in other studies^78^ and highlights the need for alternative methods to assess osteogenic differentiation. Therefore, ALP staining was used to assess osteogenic differentiation in the 3D bioprinted samples (**Figure 7**).

**Figure 7.**
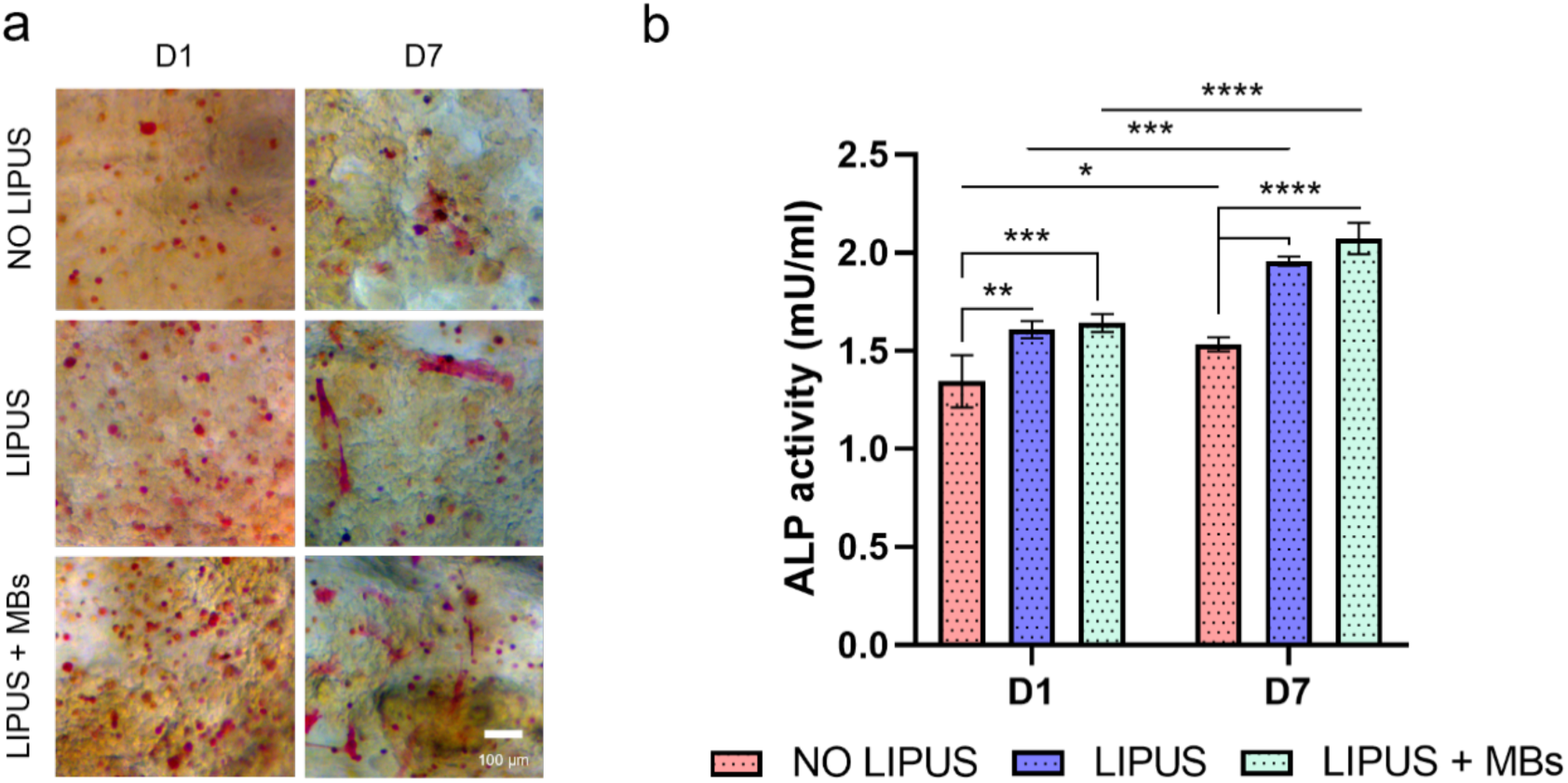
ALP activity of unstimulated, LIPUS-stimulated, and LIPUS + MBs-stimulated HBMSCs in hydrogels. a) ALP expression observed at 1-7 days after cell culture of LIPUS-stimulated (w and w/o MBs) and non-stimulated HBMSCs within scaffolds. Images were taken at 20x magnification. b) ALP activity was quantified in mU/ml. Scale bar: (a) 100 μm. Statistical significance was assessed by two-way ANOVA. Mean ± S.D. n=3, ****p<0.0001, ***p<0.001, **p<0.01, *p<0.05.

Consistently with the findings by Xia and colleagues^79^, demonstrating that the combination of LIPUS and MBs enhanced osteogenic differentiation of HBMSCs in 3D scaffolds, both LIPUS and LIPUS + MBs samples showed significantly (p<0.05) higher ALP activity than unstimulated controls on day 7. Additionally, our results also showed increased cell spreading in the LIPUS and LIPUS + MBs groups, indicating a greater physiological skeletal functionality compared to control samples. These results suggest that LIPUS stimulation effectively promoted the osteogenic differentiation of HBMSCs, with MBs further enhancing this effect by amplifying the mechanical effects of LIPUS. Immunofluorescence investigation was carried out to examine the expression of OCN and OPN in both stimulated and unstimulated samples (**Figure 8a**).

**Figure 8.**
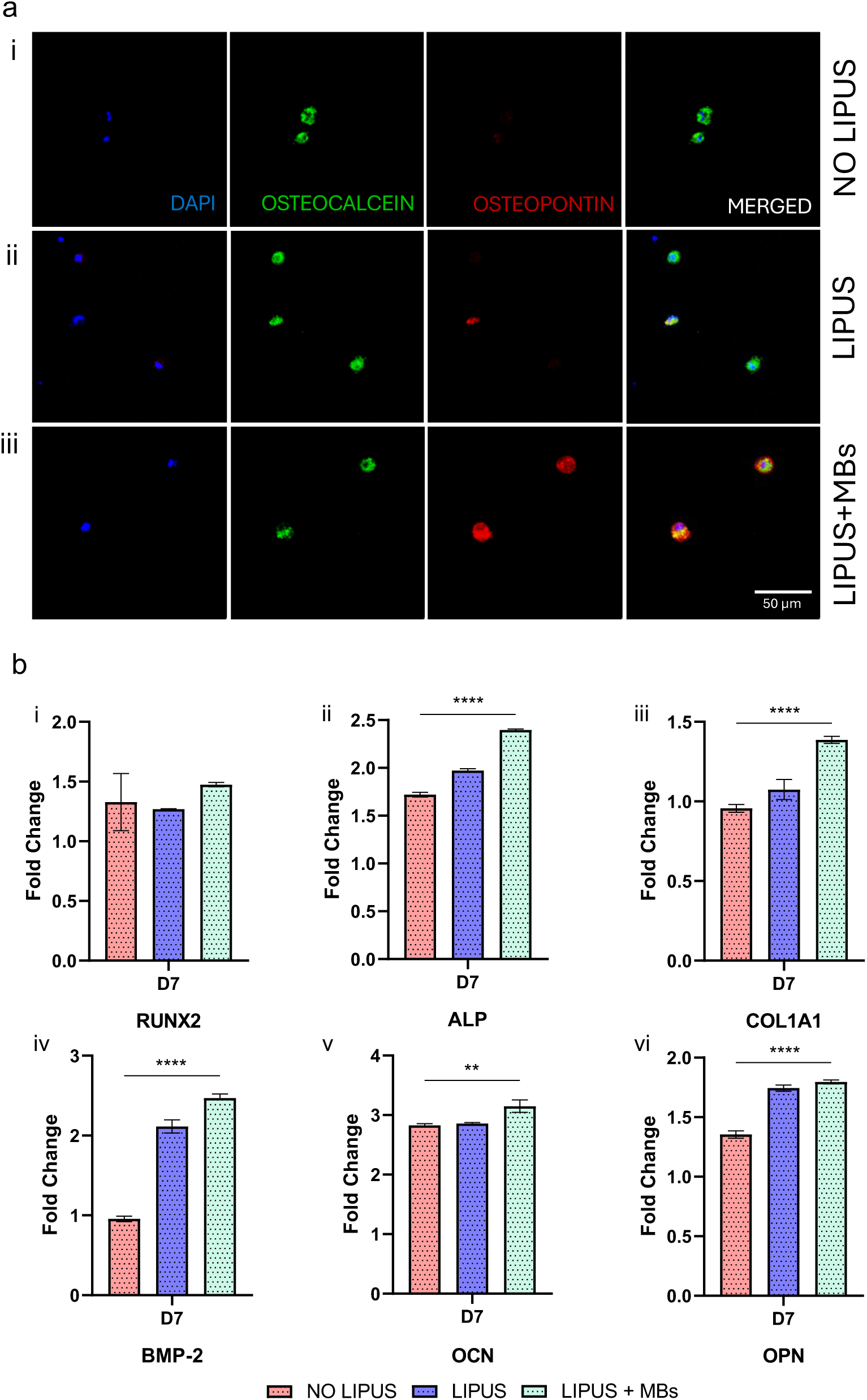
Immunofluorescence images (a) of (i) unstimulated, (ii) LIPUS-treated and (iii) LIPUS + MBs-treated HBMSCs in hydrogels after 7 days of 2D culture showing the nucleus (blue), OCN (green) and OPN (red) at 20x magnification. b) RT-qPCR analysis of HBMSCs after 7 days of 2D culture. The expression of genes associated with osteogenic differentiation was examined to observe the effect of LIPUS stimulation. The expression levels of (i) RUNX-2, (ii) ALP, (iii) COL1A1, (iv) BMP-2, (v) OCN and (vi) OPN were significantly higher in samples exposed to LIPUS at day 7 compared to controls. Scale bars: (a) 50 μm. Statistical significance was assessed by one-way ANOVA. Mean ± S.D. n=3, ****p < 0.0001, **p<0.01.

On day 7, OCN expression was significantly higher in LIPUS and LIPUS + MBs samples compared to controls. The OPN level, although less pronounced than OCN, was found to be over-expressed in LIPUS + MBs samples followed by LIPUS-only samples, with nearly non-detectable levels observed in control groups. This observation suggests an enhanced differentiation of skeletal stem cells towards more mature osteoblastic cells. These findings are consistent with previous studies reporting increased expression of these markers samples exposed to MBs and LIPUS compared to LIPUS alone^29^, further supporting the role of MBs in enhancing the osteogenic effects of LIPUS. To complement immunofluorescence with more quantitative analysis and to further assess gene expression related to osteogenic differentiation, RT-qPCR analysis was performed on samples stimulated with LIPUS, both with and without MBs (**Figure 8b**). Unstimulated samples were used as controls. By day 7, osteogenic gene expression was upregulated, including RUNX-2, ALP, COL1A1, BMP-2, OCN and OPN, with overall significantly higher expression in LIPUS + MBs samples compared to LIPUS alone. Xu and co-workers^80^ also showed that the presence of MBs significantly increased osteogenic gene expression, further supporting the hypothesis that MBs enhance LIPUS-induced gene expression, which is critical for osteoblast activity and bone formation.

In summary, these results demonstrate for the first time the application of the 3DBonUS methodology: a microfluidic-assisted 3D bioprinting system integrated with LIPUS to fabricate functional constructs promoting enhanced osteogenic differentiation. The 3D constructs fabricated via 3DBonUS hold promise as skeletal replacements to stimulate bone regeneration. Future *in vivo* studies will be required to evaluate the safety and efficacy of the 3DBonUS constructs in promoting bone regeneration and repair.

## 4. Conclusions

This study demonstrates the potential of the 3DBonUS approach, integrating LIPUS with microfluidic-assisted 3D bioprinting, in driving and enhancing the osteogenic differentiation of HBMSCs. By incorporating LIPUS stimulation directly into the 3D bioprinting process, we observed a significant increase in the differentiation of HBMSCs, supporting the creation of functional and implantable bone tissue scaffolds. The 3DBonUS platform addresses the ongoing need for improved bone regeneration techniques, by providing a non-invasive and controlled means of stimulating cells during the early stages of scaffold formation. The addition of MBs further enhanced cellular responses, as evidenced by increased ALP activity and upregulation of critical osteogenic markers including RUNX-2, ALP, COL1A1, BMP-2, OCN and OPN. The combined application of LIPUS and MBs not only accelerated osteogenesis but also enhanced bone matrix deposition, highlighting its potential to improve both the quality and speed of bone tissue regeneration. Moreover, the dual-chip microfluidic system developed in this study allowed repeatable LIPUS stimulation throughout the cell suspension during 3D bioprinting. This design approach ensured that each flowing cellular component received continuous mechanical stimulation, maximising the osteogenic potential of the entire cell population. Unlike traditional US-mediated approaches that rely on repeated long-term stimulation, the 3DBonUS approach used a single controlled LIPUS exposure during 3D deposition, driving the differentiation process while maintaining high 3D bioprinting efficiency. Future research will focus on refining the 3DBonUS printhead into a single monolithic microfluidic extrusion chip. This advancement would allow simultaneous stimulation of both the biomaterial scaffold and the embedded cells, allowing further investigation of the combined effects of LIPUS on cellular differentiation and biomaterial interactions. Such developments hold great promise for improving bone TERM applications, paving the way for more efficient and scalable methods to engineer functional bone implants for clinical use. Furthermore, *in vivo* studies will be performed in the future to validate the therapeutic efficacy of the 3DBonUS constructs for skeletal repair in more clinically-relevant scenarios.

## 5. Acknowledgements

GC acknowledges funding from MTF Biologics (OSTEOMIMIC). This research was partially funded by grants from ERC-2019-Synergy Grant (ASTRA, n. 855923); EIC-2022-PathfinderOpen (ivBM-4PAP, n. 101098989); Project “National Center for Gene Therapy and Drugs based on RNA Technology” (CN00000041) financed by NextGeneration EU PNRR MUR—M4C2—Action 1.4—Call "Potenziamento strutture di ricerca e creazione di “campioni nazionali di R&S” (CUP J33C22001130001). The authors wish to thank the microscopy facility at Center for Life Nano- and Neuro-Science, Fondazione Istituto Italiano di Tecnologia. Cartoons in figures were created with BioRender.

---

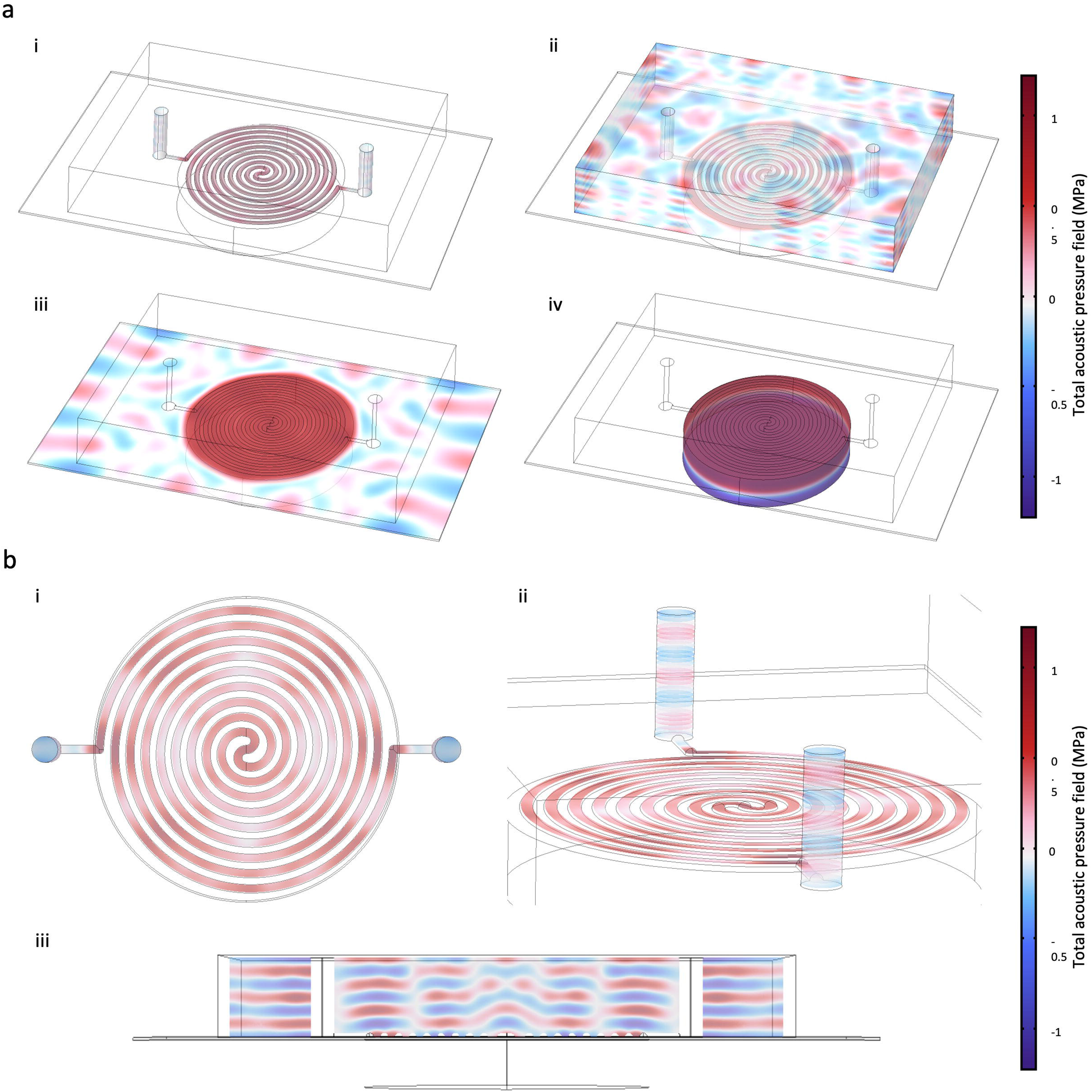

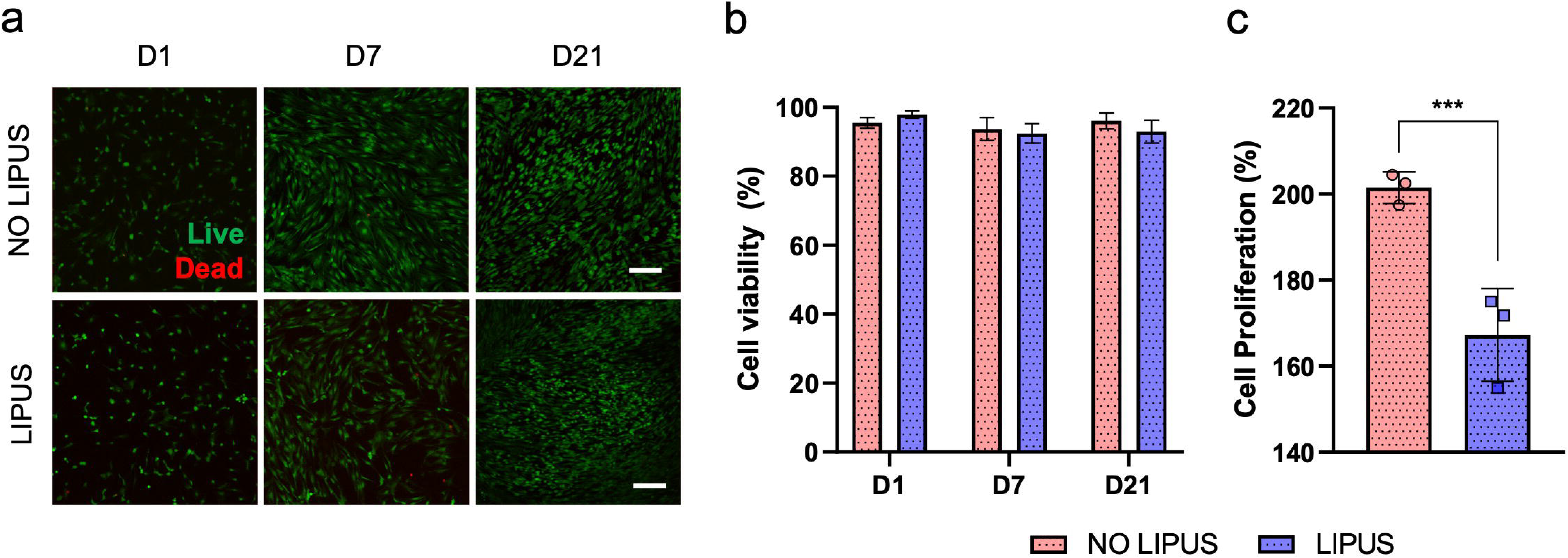

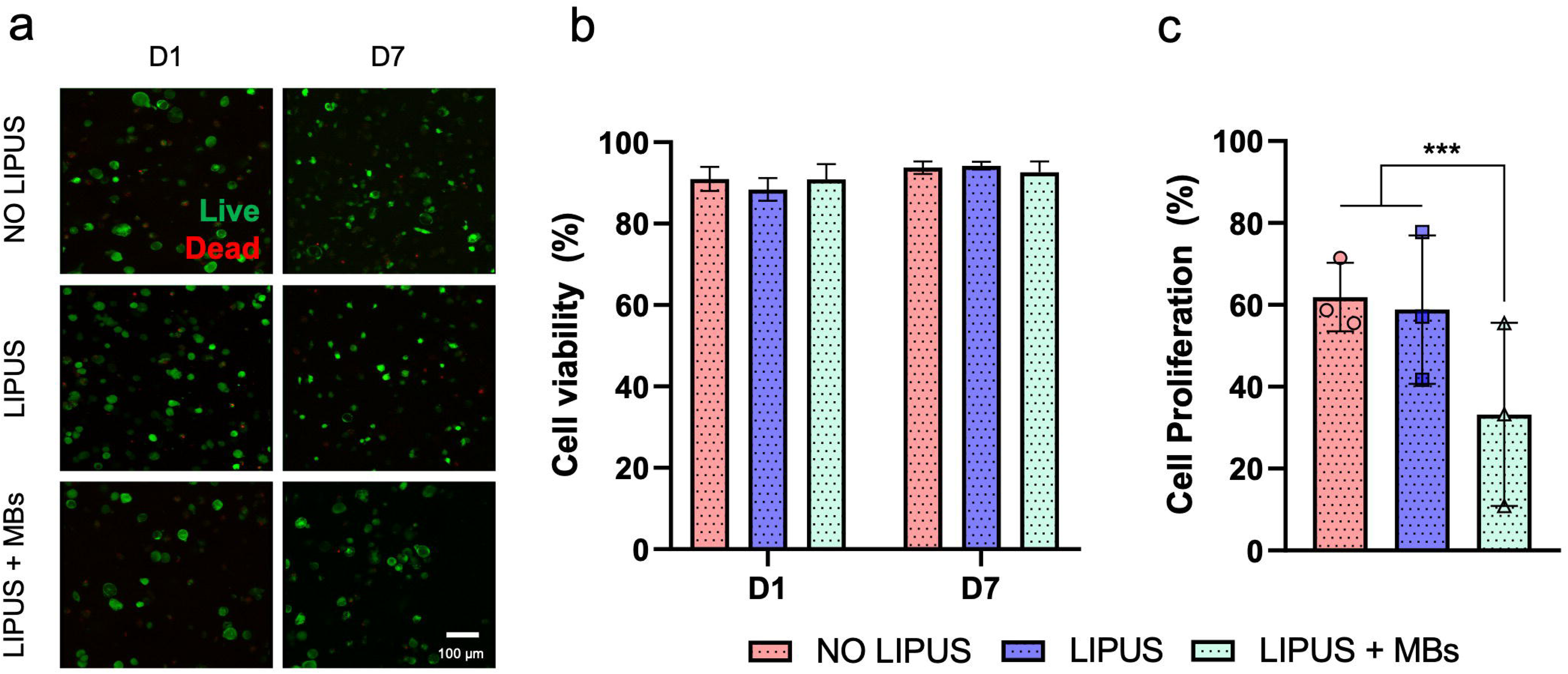

