## Supplementary Info for "3D bone printing via primed differentiation of stem cells with ultrasound (3DBonUS)"

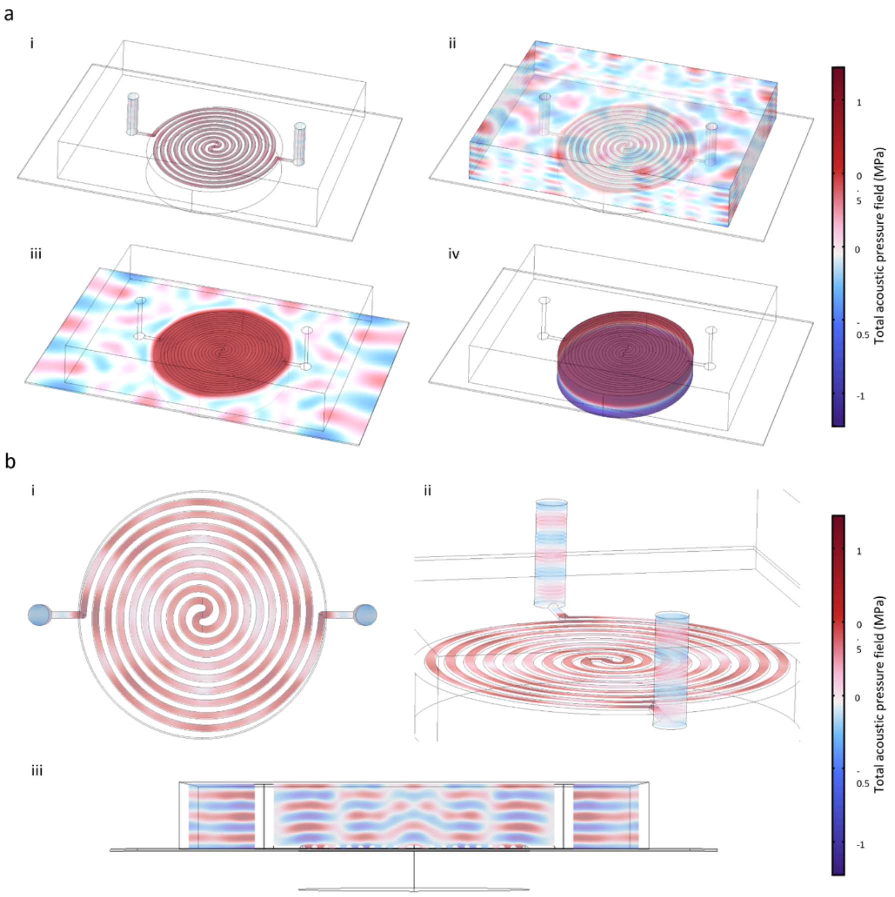


**Figure S1**. COMSOL simulations performed on the acoustic device. a) Simulated total acoustic pressure field in the a-i) spiral microfluidic channel, a-ii) PDMS chip, a-iii) 0.17 mm thick glass slide and a-iv) 647 kHz PZT disc. b) Detail of the simulated total acoustic pressure field in the spiral microfluidic channel b-i) seen from the top (one slice), b-ii) from the top (slices) and b-iii) from the PDMS chip side (slice).

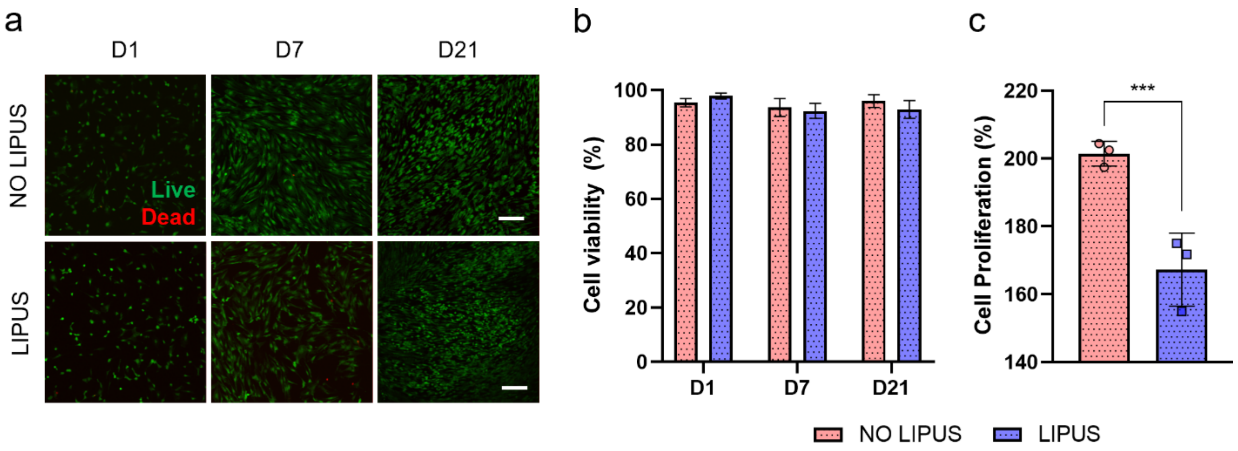


**Figure S2**. Viability and growth of unstimulated and LIPUS-stimulated HBMSCs. a) Photomicrographs of HBMSCs in 2D cultures not exposed and exposed to LIPUS on days 1-7-21. Cells were stained with Calcein to observe live cells (green) and propidium iodide to observe dead cells (red). b) Quantified cell density of live and dead HBMSCs. c) Percentage of cell growth calculated on day 21 of cell culture. Scale bar: (a) 200 µm. Mean ± S.D. n=3. ***p<0.001.

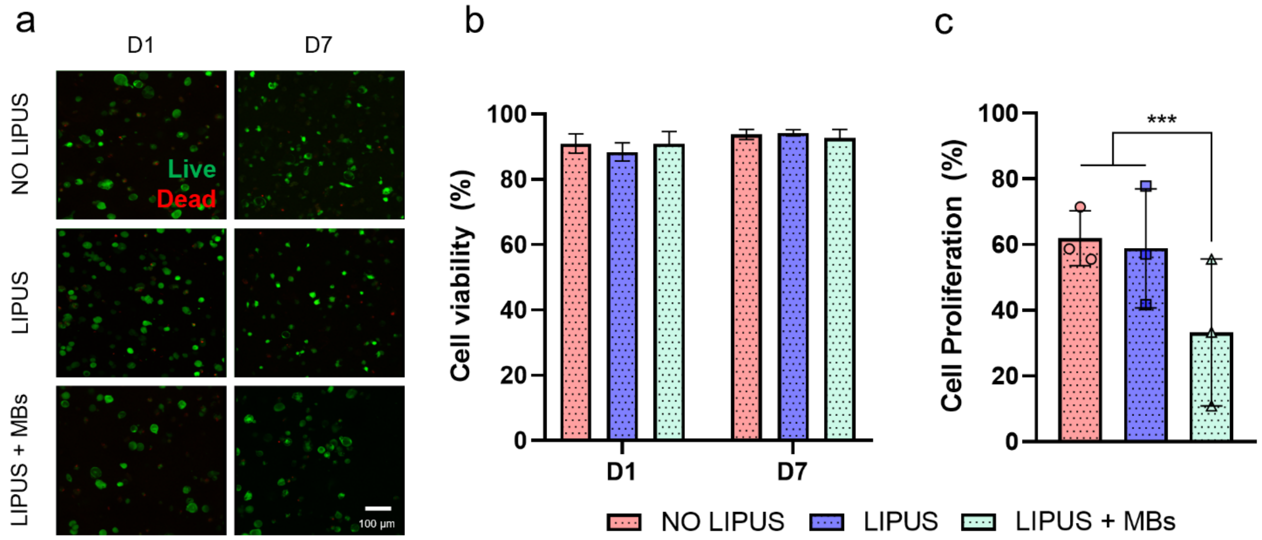


**Figure S3**. Cell viability and growth of unstimulated, LIPUS-stimulated and LIPUS + MBs-stimulated HBMSCs in hydrogels. a) Photomicrographs of HBMSCs in 3D cultures not exposed to LIPUS, exposed to LIPUS, and exposed to LIPUS + MBs on days 1-7. Cells were stained with Calcein to observe live cells (green) and propidium iodide to observe dead cells (red). b) Quantified cell density of live and dead HBMSCs in GelMA + AlgMA hydrogels. c) Percentage of cell growth calculated on day 7 of cell culture. Scale bar: (a) 100 µm. Mean ± S.D. n=3. ***p<0.001.
